# *In vivo* cartography of state-dependent signal flow hierarchy in the human cerebral cortex

**DOI:** 10.1101/2025.06.24.660962

**Authors:** Younghyun Oh, Yejin Ann, Jae-Joong Lee, Takuya Ito, Sean Froudist-Walsh, Casey Paquola, Michael Milham, R. Nathan Spreng, Daniel Margulies, Boris Bernhardt, Choong-Wan Woo, Seok-Jun Hong

**Affiliations:** Center for Neuroscience Imaging Research, Institute for Basic Science, Suwon, South Korea; Department of Biomedical Engineering, Sungkyunkwan University, Suwon, South Korea; Life-inspired Neural Network for Prediction and Optimization Research Group, Suwon, South Korea; Department of Intelligent Precision Healthcare Convergence, Sungkyunkwan University, Suwon, South Korea; T.J. Watson Research Center, IBM Research, Yorktown, NY, United States; Bristol Computational Neuroscience Unit, School of Engineering Mathematics & Technology, University of Bristol, Bristol, United Kingdom; Institute of Neuroscience and Medicine, Forschungszentrum Jülich, Germany; Nathan S. Kline Institute for Psychiatric Research; Center for the Developing Brain, Child Mind Institute, New York, NY, United States; McConnell Brain Imaging Centre, Montreal Neurological Institute and Hospital, McGill University, Montreal, QC; Integrative Neuroscience and Cognition Center, CNRS, Université de Paris Cité, Paris, France; Department of MetaBioHealth, Sungkyunkwan University, Suwon, South Korea

## Abstract

Understanding the principle of information flow across distributed brain networks is of paramount importance in neuroscience. Here, we introduce a novel neuroimaging framework, leveraging integrated effective connectivity (iEC) and unconstrained signal flow mapping for data-driven discovery of the human cerebral functional hierarchy. Simulation and empirical validation demonstrated the high fidelity of iEC in recovering connectome directionality and its potential relationship with histologically defined feedforward and feedback pathways. Notably, the iEC-derived hierarchy revealed a monotonically increasing level along the axis where the sensorimotor, association, and paralimbic areas are sequentially ordered – a pattern supported by the Structural Model of laminar connectivity. This hierarchy was further demonstrated to flexibly reorganize across brain states: flattening during an externally oriented condition, evidenced by a reduced slope in the hierarchy, and steepening during an internally focused condition, reflecting heightened engagement of interoceptive regions. Our study highlights the unique role of macroscale directed functional connectivity in uncovering a biologically interpretable state-dependent signal flow hierarchy.

## Introduction

Human cognition arises from ever-changing dynamical flows of neural information across distributed brain networks. Cortical hierarchy, one of the most fundamental architectures for this functional dynamics, has been extensively investigated in previous studies, with a focus on the computational principles of sensorimotor hierarchies in early work^1–7^, while recently more expanded to the whole-brain level^8–10^ to highlight its global basis involved in a wide range of cognitive functions. Indeed, a rich array of evidence from histology^11,12^, tract-tracing^13–15^, and functional MRI (fMRI)^16–18^ studies has shown that both human and non-human primate brains exhibit a unique large-scale organizational axis spanning from the sensorimotor to association cortices, termed as ‘functional gradient’^9^ or ‘global processing hierarchy’^9,19^. Notably, this macro-scale axis has been reproduced across many neuroimaging studies^20–25^ and related to multiple cognitive domains such as perception, action and social processing^24,26–28^, typical development^22,29,30^ and system-level pathogenicity in neuropsychiatric conditions^17,31,32^. Despite its significance, exactly how the hierarchical organization emerges from dynamical neural information flow remains poorly understood^9,33^.

Two major issues contribute to this lack of understanding: 1) the state-dependent organization of functional brain dynamics and 2) the limitations of conventional fMRI connectome analytics. The former arises from the brain’s frequent state transitions. Indeed, in daily life, our brain continuously shifts between different states, driven by exteroceptive stimuli, such as watching a movie with a high influx of multisensory information, as well as interoceptive signals, such as hunger or thirst reflecting the ongoing physiological states of the body. Even at rest, the brain dynamically organizes neural pathways to generate diverse thoughts from continuous mind wandering. While this “*state-dependent functional dynamic*” is a fundamental characteristic of the brain for adaptive behaviors, so far only few studies have systematically compared different brain states with respect to their dynamics, especially in the context of information flow along the cortical hierarchy.

The second issue is that the current network neuroscience approach predominantly relies on *undirected* functional connectivity (FC). Given that the brain network is a constellation of feedforward and feedback pathways, the inability to infer connectome directionality significantly impedes the field’s advancement. This challenge has spurred the development of an alternative approach—*effective connectivity* (EC) mapping—a ‘directed version’ of connectome analytics. This technique, which enables the statistical inference of the net influence of one brain area on another, has been increasingly sophisticated and improved in recent studies, now supporting even the whole-brain assessment of directed connections based on fMRI data^34–36^. As these methods flourish, however, EC research is encountering new challenges related to algorithm validation and selection^37^, which are crucial to address to move forward to the next generation of EC approaches. These issues are summarized as follows:

*i) Overabundance of EC algorithms*. Over the last two decades, the field has witnessed the proposal of numerous EC algorithms^35,36,38–49^, each based on a unique mathematical principle. This diversity, while testament to the field’s vibrant interest, paradoxically resulted in a lack of consensus, which made it unclear when and how to use each algorithm and raised the concerns for reproducibility of findings^40,48,50–54^, as different methodological foundations may lead to unintended variations in the results.
*ii) Network complexity*. While different EC methods have unique strengths in their statistical properties, most of them have not been sufficiently tested for high-complexity network data (*e.g.,* high-resolution network nodes with dense cyclic and negative connections), and except for few recent studies^34,35^, their scale often remained on the order of several dozen nodes^40,47,55–58^. As shown in recent approaches of brain parcellation, however, the human cerebral cortex can be divided into at least ≥100 functionally and/or structurally distinct areas^59,60^. Developing EC algorithms that are scalable to networks with high dimensionality and complexity is therefore imperative to precisely infer the functional dynamics in the human brain.
*iii) Lack of biological validation*. EC algorithms are typically validated using artificially simulated networks rather than biological data. This is particularly true in the human brain, where ground truth remains unknown. This lack of biological validation often leads to a limited use of EC analyses, mainly for those studies having a clear hypothesis on the connectivity configuration of a targeted circuit^61–63^ (but see recent advances^64–66)^.

Here, we propose a unified imaging analysis framework termed ‘*integrated EC (iEC)*’ to address these challenges. This approach is based on the idea of combining multiple existing EC algorithms with distinct mathematical properties^37^ in order to synergize their complementary strengths while obviating the development of yet another statistical algorithm. To implement this, we underwent an extensive validation of individual EC algorithms and integrated them into the iEC framework, testing on the simulation of biologically plausible networks and comparing with the empirical tract-tracing and resting-state fMRI data across two different network complexities, both comprising ≥100 nodes and dense cyclic connections.

After demonstrating the robustness of our approach across three independent validations, we used this iEC framework to thoroughly chart out the characteristics of directed functional connectivity in the human brain. This analysis revealed distinct connectional polarity along the cortical hierarchy, with unimodal sensory areas predominantly exhibiting positive (strengthening) outgoing connections, whereas heteromodal association areas exhibit a balanced ratio of positive and negative (suppressing) outgoing connections. Given that these patterns closely mirror converging evidence on the biological characteristics of feedback and feedforward connections in previous studies^67–69^, we further leveraged a data-driven method to estimate a cortex-wide hierarchical level based on iEC. This provided, for the first time, a macroscale map of ‘*signal flow hierarchy’* in the human brain. Indeed, this map showed a remarkably similar pattern to a previously conceptualized brain hierarchy model (*i.e.,* Mesulam’s proposition^19^) but also to cytoarchitecturally determined cortical types^70^, both spanning from low-level sensory (koniocortical) to higher-order (eulaminate and dysgranular) areas and up to the paralimbic (agranular) regions, the latter mainly known for interoceptive cortical areas that relay signals from the ‘internal milieu’ (*i.e.* body and organs)^71^. Lastly, we demonstrated that the identified cortical hierarchy undergoes considerable state-dependent reorganization by comparing the hierarchy map of the resting state to those of two other (non-resting) brain states, *i)* a movie-watching condition that mainly evokes externally oriented process, and *ii)* a tonic pain condition (induced by oral capsaicin), which evokes more interoceptive bodily sensation. Together, our iEC framework offers a unique opportunity for *in-vivo* cartography of directed functional hierarchy in the human brain and paves the way toward a quantitative delineation of state-dependent network information flow.

## Results

Our study consists of two main phases. First, we examined the distinct characteristics of individual EC algorithms. While investigating the strengths and weaknesses of each method, we developed a new approach to integrate their results (integrated EC, hereafter called ‘iEC’), which demonstrated significantly improved accuracy of EC mapping through a multi-step validation process. In the second phase, we utilized this iEC to comprehensively examine the human brain in terms of i) global patterns of unconstrained signal flows, ii) retrieval of signal flow hierarchy, and the state-dependent reorganization of this hierarchical structure.

**Figure 1.**
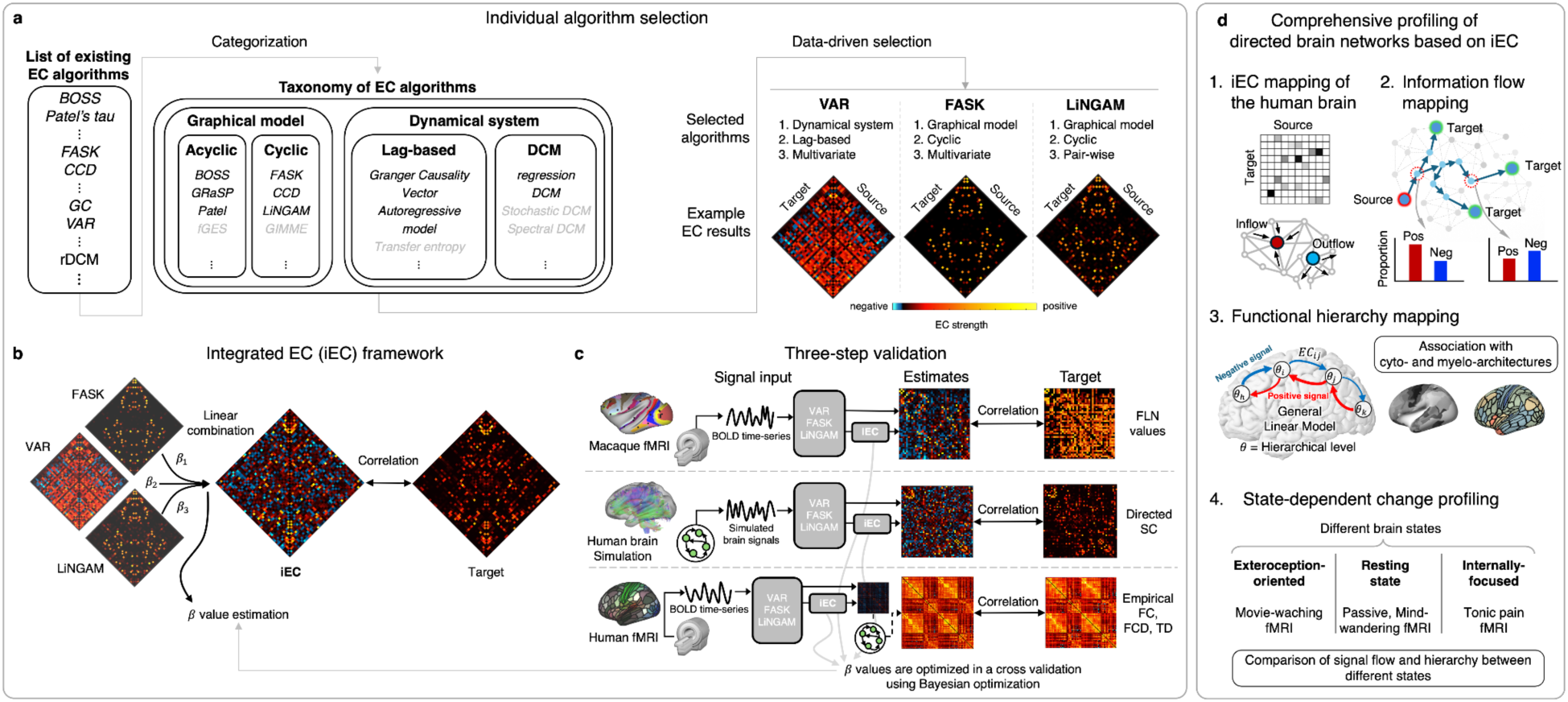
Overview of the iEC approach and comprehensive network profiling of the human brain. **a**, Taxonomy of EC algorithms. After comprehensive research and classification, we have categorized individual EC algorithms into two major groups, one based on “*Graphical models*” and the other based on “*Dynamical systems*”. Among these, we selected representative EC algorithms for each category through a fully data-driven validation, which included VAR, FASK and LiNGAM. **b**, The results of individual algorithms were then combined using Bayes-optimized linear integration, which performs a weighted sum of their EC results to create an iEC. **c**, Both individual and iEC algorithms were evaluated through a three-step validation process: *i)* empirical fMRI signals from the macaque brain were fed into each EC algorithm and the results were validated against tract-tracing based ‘Fraction of Labeled Neuron’ (FLN); *ii)* simulated brain-like signals were input into each algorithm and the estimated EC was compared to ground-truth synthetic directed structural connectivity in the human brain; *iii)* empirical fMRI signals from the human brain were input into each algorithm, the resulting EC matrix was used to simulate brain signals, and the simulated FC (functional connectivity), FCD (dynamics), and TD (time delay) were compared against empirical data **d**, After methodological validation, the group-level iEC map from empirical human fMRI was profiled in terms of (1) degree distribution and (2) the signal flow across the network. Next, (3) the hierarchical levels across the whole brain were estimated using a general linear model based on the iEC values and evaluated based on architectonic data, and finally (4) compared across different brain states including externally-oriented, resting and internally-oriented states.

To develop our iEC framework, we included widely used 9 following individual algorithms, each previously demonstrated to yield high EC mapping accuracy: regression Dynamic Causal Modeling (rDCM^35^), Vector Autoregressive model (VAR^72^; a lag order of 1), Multivariate Granger Causality (MVGC^73^), Fast Adjacency Skewness algorithm (FASK^40^), Causal Cyclic Discovery (CCD^74^), Best Order Score Search algorithm (BOSS^34^), direct linear non-Gaussian acyclic model (directLiNGAM^75^), Greedy Relaxations of the Sparsest Permutation algorithm (GRaSP^76^), and Patel’s Tau^77^ (see **Table 1** in **Methods** for details). Although non-exhaustive, our inclusion intended to cover as a wide spectrum of methodological principles as we could, targeting both linear/nonlinear, pairwise/multivariate, and Graphical/Dynamical systems algorithms among those comprehensively benchmarked in the prior studies (**Supplementary Fig. 1**). These chosen algorithms were then integrated to construct the iEC. To this end, we computed a weighted sum of individual EC results, in which the weight coefficient was optimized to yield the iEC showing the highest correlation with the target connectivity metrics (*e.g.,* ground-truth directed networks). To prevent overfitting, we optimized the weights in a strict cross-validated manner, employing Bayesian optimization (see **Methods** ‘*2.3 Integrated EC framework*’).

**Table 1.** Summary of effective connectivity (EC) algorithms used in the iEC framework. Each algorithm is categorized by its primary modeling class, methodological subtype, whether it assumes linear or nonlinear interactions, and whether it operates in a multivariate or pairwise manner.

| Method | 1st Category | 2nd Category | Linearity | Scope |
| --- | --- | --- | --- | --- |
| rDCM | Dynamical systems | Dynamic causal models | Linear | Multivariate |
| VAR | Dynamical systems | Lag-based | Linear | Multivariate |
| MVGC | Dynamical systems | Lag-based | Linear | Multivariate |
| CCD | Graphical model | Hybrid | Nonlinear | Multivariate |
| FASK | Graphical model | Hybrid | Linear | Multivariate |
| BOSS | Graphical model | Bayes Net | Linear | Multivariate |
| GRaSP | Graphical model | Bayes Net | Linear | Multivariate |
| DirectLiNGAM | Graphical model | Hybrid | Linear | Pairwise |
| Patel's tau | Graphical model | Hybrid | Nonlinear | Pairwise |

### 1. Three-step validation of EC algorithms

To comprehensively evaluate the accuracy of individual algorithms and iEC, we conducted a multi-step validation procedure. Specifically, it comprised: *(i)* comparison against empirically derived directed anatomical connectivity from the macaque brain via viral tracer injections; *(ii)* simulations utilizing biologically plausible directed networks constructed from the diffusion MRI-based structural connectivity; and *(iii)* assessment of EC algorithms’ capability to reproduce dynamics inherent to empirical fMRI signals of the human brain. The details are as follows.

#### 1.1 Validation using macaque tract-tracing and fMRI data

We began our validation by leveraging macaque tract-tracing^10^ and resting-state fMRI data (PRIME-DE^78^), recognizing the macaque resource as a gold-standard reference for directed whole-brain connectivity in the primate species (**Fig. 2a**). Individual EC algorithms were applied to resting-state fMRI from 19 anesthetized macaques, parcellated using the M132 atlas^13^. Using a randomly selected training subset (N=9), we estimated optimal integration weights (β values) for each algorithm. The median βs from the training set were then used to construct iEC matrices in the test set (N=10). Validation was performed by comparing the resulting EC estimates against “Fraction of Labeled Neurons” (FLN) data, a tract-tracing metric that quantifies directed anatomical connectivity via histological labeling (see **Methods** ‘*1.1 Macaque tract-tracing data*’ for details).

**Figure 2.**
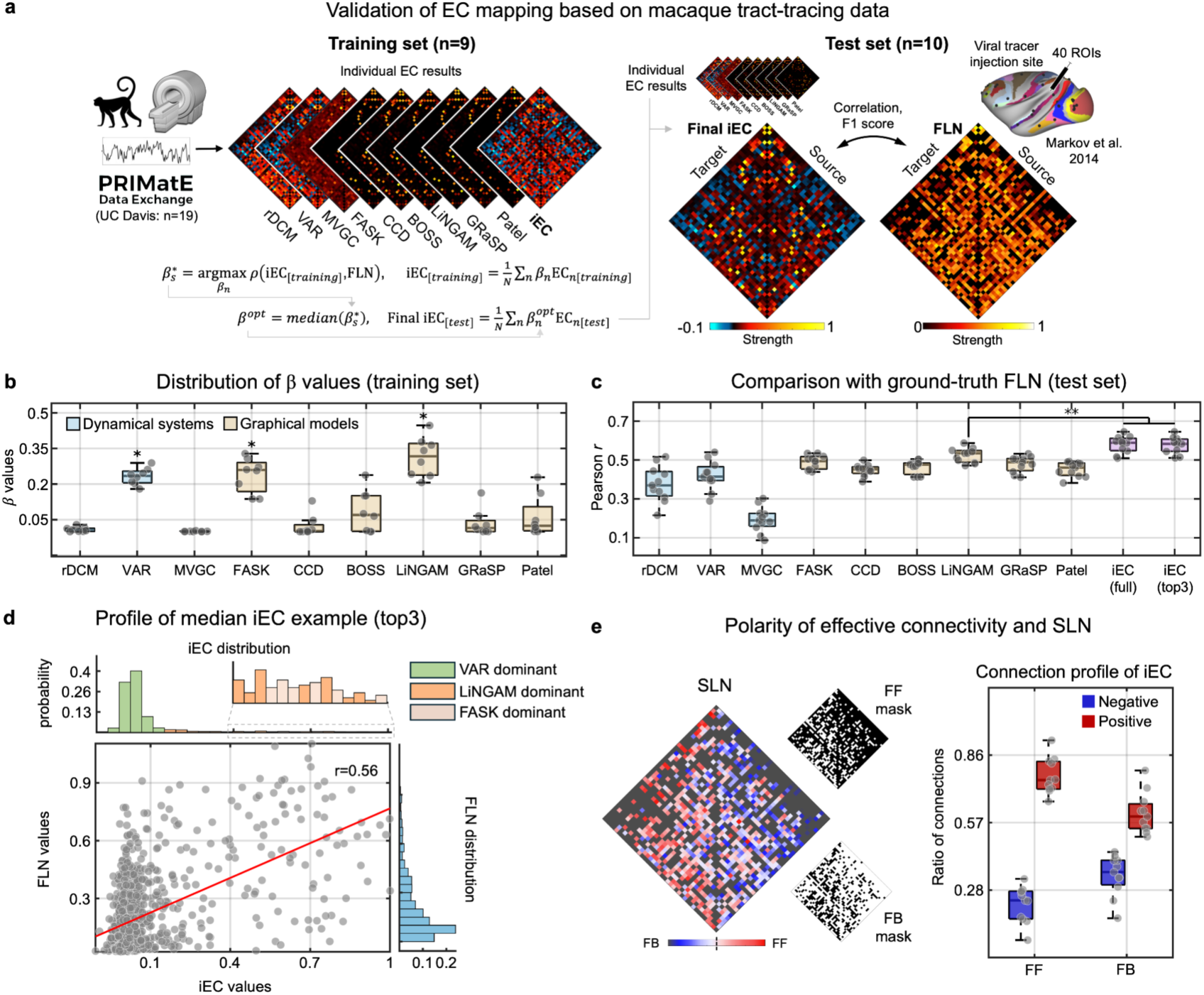
Validation of individual and iEC algorithms using macaque tract-tracing data. **a**, Individual ECs were inferred from anesthetized macaque fMRI data, which was parcellated into 40 regions defined by viral tracer injection sites. Optimal β coefficients for each EC algorithm were derived from the training dataset (N = 9) by maximizing the correlation between integrated EC (iEC) and the fraction of labeled neurons (FLN). Using median optimal βs, we constructed iEC from an independent test dataset (N = 10) to assess generalizability. **b**, Box plot of optimal βs derived from the training dataset. Boxes are color-coded by methodological families: dynamical systems (blue) and graphical models (yellow). Three algorithms—VAR, FASK, and LiNGAM—demonstrated notably higher β values (p<0.001 in one-tailed t-test, FDR-corrected), indicative of significant contribution. **c**, Performance comparison of individual EC and iEC algorithms. Two iEC variants were evaluated: one integrating all 9 algorithms and another using only the three algorithms with statistically significant β values. **d**, Analysis interpreting the enhanced performance of iEC. Data represent a single macaque brain with median performance for iEC (top three), showing a correlation of 0.56 with FLN. The FLN distribution exhibited a strong heavy-tail structure (right vertical histogram), with the tail primarily captured by LiNGAM and even more so by FASK (upper landscape histogram). In contrast, VAR primarily captured weaker connections, which, despite their relative weakness, accounted for more than 50% of the total connections. **e**, Connection polarity investigated via supragranular labeled neurons (SLN). Connections were categorized into feedforward (FF) and feedback (FB) pathways based on an unbiased SLN threshold of 0.5, and iEC values were profiled separately for each pathway.

EC algorithms revealed varying degrees of correlation with the ground truth of FLN data (**Fig. 2**). Notably, their optimized β values showed a dominant contribution in only one or two algorithms per methodological family (**Fig. 2b**): VAR (dynamical systems), LiNGAM and FASK (both graphical models) (one-tailed t-test, p<0.001, FDR-corrected). Based on these findings, we constructed two iEC variants—one using all algorithms (full iEC) and another using the top three (VAR, FASK, LiNGAM; top3 iEC)—and estimated their performance using test macaque subjects (**Fig. 2c**). Both variants yielded similar accuracies (median r=0.60) while significantly outperforming the best-performing individual algorithm (median r=0.52; p<0.001, Wilcoxon signed-rank test). To complement this correlation-based metric, we further evaluated directionality of identified ECs using an F1 score. After binarizing the EC matrices at two sparsity thresholds (15%, 30%), we computed an F1 score based on precision and recall against FLN. Again, iEC outperformed individual algorithms at both thresholds (p=0.02 at 15%, p<0.001 at 30%; **Supplementary Fig. 2**), confirming its superior accuracy in directionality mapping.

To investigate the source of this superiority, we scrutinized the contribution of the three individual algorithms used to construct iEC (**Fig. 2d**). It should be noted that FLN exhibits a heavy-tailed distribution typical of biological systems, indicating a non-trivial proportion of the strong connections^79,80^. Interestingly, each algorithm was found to specialize in a different regime of this distribution axis. For instance, VAR captured widespread, weak-strength connections, while LiNGAM and FASK targeted the detection of sparse yet strong connections. FASK was found to specialize in particularly the upper extreme, *i.e.,* the strongest ECs (**Supplementary Fig. 3**). These results highlight the complementary role of each algorithm in reconstructing the EC landscape.

We applied these algorithms directly to Blood Oxygenation Level Dependent (BOLD) signals, which are typically known to represent hemodynamic responses (HR) mediated by underlying neurovascular coupling. To investigate the influence of these HR on our EC estimation, we re-tested all individual and iEC algorithms by applying a state-of-the-art brain-area-specific HR deconvolution method to empirical macaque BOLD signals (see **Methods** for details). Most algorithms, with the exception of MVGC, revealed strikingly high consistency of EC maps between pre- and post-deconvolution signals (all individual methods: r>0.8). This was especially true for the iEC approach (r=0.91; see **Supplementary Fig. 4** for the replication of this result in the human brain [r=0.87]). MVGC was excluded from all subsequent analyses due to its low consistency for HR effects. Based on these largely intact EC results, together with prior work showing rather compromised system identification after HR deconvolution^81^, we proceeded with the subsequent EC mapping analyses using the original BOLD signals.

Importantly, the visual inspection of iEC results (**Fig. 2a**) indicated that positive connections (red color-coded) are both stronger and more prevalent, while negative connections (blue color-coded) across the brain areas are generally weak and sparse. Previous studies suggested that these connection polarities may reflect intricately mixed excitatory and inhibitory effects^82,83^, potentially aligned with feedforward (FF) and feedback (FB) signal pathways^67^. Indeed, while a dominant portion of FF connections have been previously found to target excitatory neurons (thus exerting positive and amplifying influences to target regions)^84^, FB connections were associated with both excitatory and inhibitory pathway^85–87^, although the later, ‘*suppressive*’ effect has traditionally been considered a major characteristic of FB. To assess whether these distinct patterns can be also captured in our macro-scale analysis, we profiled the iEC with respect to SLN (Fraction of Supragranular Labeled Neurons), a marker to quantitatively index the level of FF and FB connectivity (see **Methods** ‘*1.1 Macaque tract-tracing data*’). Interestingly, when we categorized iECs into FF or FB by applying 0.5 threshold to their SLN value (range: 0-1; FF: SLN>0.5; FB: SLN<0.5), the majority of iECs corresponding to the FF pathway were positive, while those corresponding to FB showed a relatively more balanced ratio of positive and negative connections (**Fig. 2e** right), mirroring previous reports on signal influences exerted by FF/FB pathways.

#### 1.2 Validation using directed SC networks

While our initial validation using macaque FLN confirmed that the iEC framework can reliably capture experimentally derived ground truth, its applicability to the human brain remains uncertain, primarily due to the lack of an equivalent whole-brain directed connectivity reference. To address this challenge, we devised a carefully curated simulation approach based on the empirically derived structural connectivity (SC) from diffusion MRI data. We utilized the group-averaged SC matrices at two different parcellations (Schaefer-100^59^ and MMP-360^60^) to assess the impact of network complexity. Directionality was introduced by minimally rewiring the edges such that their directionalities were induced while preserving key topological features of the original network, including degree distribution, clustering coefficients, and global connectedness. (**Supplementary Fig. 5**). Based on these directed SC templates, we simulated whole-brain neural activity using the Hopf model—a well-established dynamical framework for generating oscillatory neural signals^88–90^ (see **Methods** and **Supplementary Fig. 6** for the details of this analysis and quantitative comparison with yet another biologically detailed simulation model). We then applied the same EC estimation and cross-validation procedures used in the macaque analyses, comparing outputs to the synthetic ground truth.

Across both low- and high-resolution brain parcellations, the EC mapping demonstrated highly consistent findings with macaque results. Indeed, the distributions of optimized β values closely mirrored those from the macaque training set (**Supplementary Fig. 7a**), and the iEC consistently outperformed individual algorithms in recovering a directed-connectivity structure across two network resolutions (**Supplementary Fig. 7b**). Taken together, our simulation analysis provides robust evidence for the resilience and scalability of the iEC framework, supporting its translational value in the human brain.

#### 1.3 Validation using human fMRI

Finally, we evaluated the EC algorithms using human resting-state fMRI (from the HCP 440-subjects training/testing cases; see **Methods** *3.3 Cross validation strategy*). Given the absence of an established ground truth, we adopted a model-based validation strategy to assess how well each inferred EC matrix could reproduce empirical signal properties of the human brain (**Fig. 3a**). Specifically, we used each EC matrix as a ‘directed-connectivity scaffold’ for simulating whole-brain dynamics and compared the resulting simulated signals to empirical fMRI data. The underlying rationale for this approach was that should the inferred EC effectively approximate an inherent structure of directional influences governing whole-brain dynamics, the resulting simulated signal would closely replicate its empirical counterpart.

**Figure 3.**
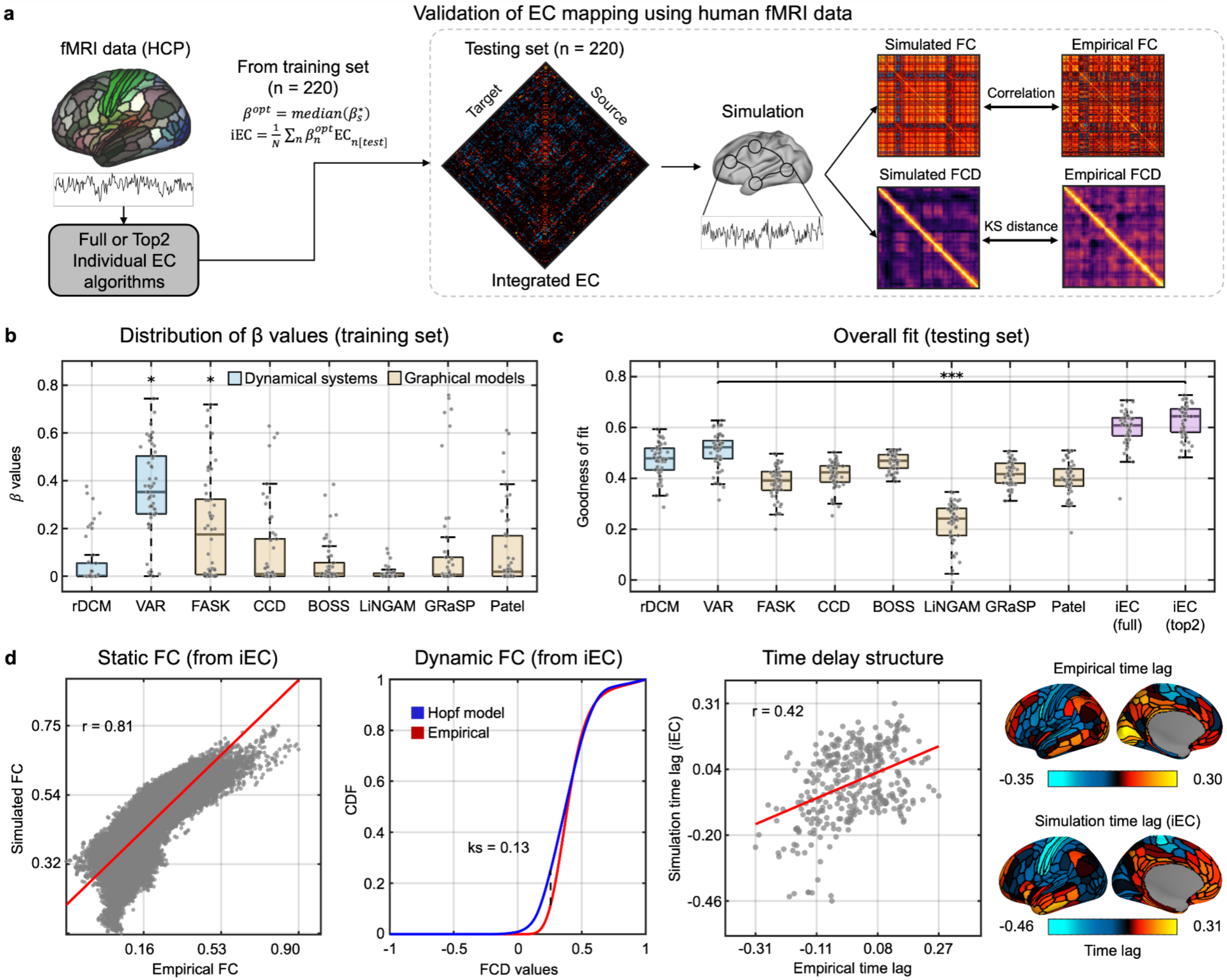
Validation of iEC framework using human fMRI data. **a**, Schematic of the validation. Individual ECs were first estimated from resting-state fMRI data. The iEC was then constructed by integrating individual ECs using the optimal β coefficients derived from a training set (N = 220). For validation, the resulting EC matrices as well as the constructed iEC were used as a network backbone to simulate fMRI signals via the Hopf model. The simulated time series were then used to compute the FC (functional connectivity) and FCD (functional connectivity dynamics), which were compared to their empirical counterparts. **b**, Distribution of the optimized βs across 50 independent runs of Bayesian optimization on the training set. Each dot represents the optimal weight obtained from a single run, reflecting each algorithm’s relative contribution to the integrated EC model. **c**, The overall fit scores of each EC algorithm and iEC evaluated in a held-out testing set (N = 220). The fit was computed as the static FC correlation minus the FCD KS distance. Each dot represents one realization of the simulation. **d**, Comparison of empirical and iEC-simulated FC/FCD. (left) Each dot represents a FC value between a pair of brain regions. The red line shows the least-squares linear fit. (middle) Cumulative distribution functions (CDFs) of FCD from empirical and simulated data, with the distance along the y-axis indicating a KS distance. (right) Comparison of empirical and simulated time delay structure. Each dot in the scatter plot represents the mean time lag of a brain region; red line denotes the best linear fit. Group-averaged delay maps are shown on the cortical surface.

To test this, we employed two complementary metrics: static functional connectivity (FC) and functional connectivity dynamics (FCD)^88,91,92^. To quantify their combined effect, we adopted a composite scoring method from previous studies^93,94^, defined as ‘Pearson correlation (*r*) between empirical and simulated FC’ minus ‘Kolmogorov–Smirnov distance (KS) between their FCD distributions’ (*i.e.,* overall fit = *r* - *KS*). A higher score signifies a closer correspondence between the simulated and empirical data’s signal properties. We leveraged this composite score to optimize the β value for each algorithm in the construction of iEC within the training dataset. Our primary analysis focused on group-level results from the MMP-360 parcellation (see **Supplementary Fig. 8** for the results using the Schaefer-100 atlas). Upon examining the optimized β values, we found that only VAR and FASK made statistically significant contributions to the iEC estimation (one-tailed t-test, p<0.001, FDR-corrected; **Fig. 3b**). This finding closely aligns with the algorithmic contributions observed in the macaque brain, thereby underscoring the robustness of our iEC construction method.

Based on this observation, we constructed two variants of iEC: one incorporating all algorithms (full iEC) and the other limited to those with significant contributions (top2-iEC), and tested their generalizability on the held-out testing set. The EC mapping results from these two models were consistent with the findings in the macaque brain (**Fig. 3c**). Specifically, both iEC variants significantly outperformed the best-performing individual algorithm (VAR). In fact, the top2-iEC yielded even a higher overall fit compared to the full iEC (Wilcoxon signed rank test, one-tailed, *p*=0.013), highlighting an advantage of implementing a parsimonious EC model. Indeed, when evaluating the two metrics comprising the fit score based on test dataset (**Figs. 3d**), the simulated FC was highly correlated with an empirical one (*r* = 0.81), and FCD distributions were also closely matched (*KS* = 0.13; the value closer to zero indicating better model fit), suggesting strong fidelity in both spatial and temporal domains. We also investigated another independent dynamics-sensitive measure called ‘time-delay (TD)’, a metric quantifying an intrinsic lag relationship across different brain regions^95^. Although TD was not included in the β optimization process, iEC-based simulations reproduced the empirical TD with a highly similar pattern (*r* = 0.42; **Fig. 3d** right).

Collectively, these multi-level assessments confirmed the validity and robustness of our iEC framework across varying network complexities and species, highlighting its capacity to capture both static and dynamic functional properties of the directed connectivity network.

### 2. Investigation of iEC profiles and signal flow hierarchy in the human brain

#### 2.1 Profiling of the human whole-brain iEC

Having thoroughly validated our iEC method, we now turn into applying this technique to comprehensively profile human whole-brain EC architectures, using a group-level iEC matrix derived from resting-state fMRI of 220 healthy young adult, test subjects.

Organizing the connectivity matrix according to the Yeo-Krienen atlas^96^ revealed a distinct connectome organization (**Fig. 4a**): connections within modules were predominantly positive, while those between modules mainly fell into the negative range, which may reflect a nature of functional organization in terms of network integration and segregation. Furthermore, in line with previous electrophysiological recording and retrograde tract-tracing experiments^79,80^, the ECs in the human brain exhibited a heavy-tailed distribution of connectivity strength (tail index=1.45; the index of <2 indicates heavy-tailedness of a distribution^97^), indicating the presence of infrequent yet strong positive connections (**Fig. 4b**). In contrast, the negative connections showed an opposite pattern, characterized by much weaker strength but with a considerable proportion in the entire network (40%).

**Figure 4.**
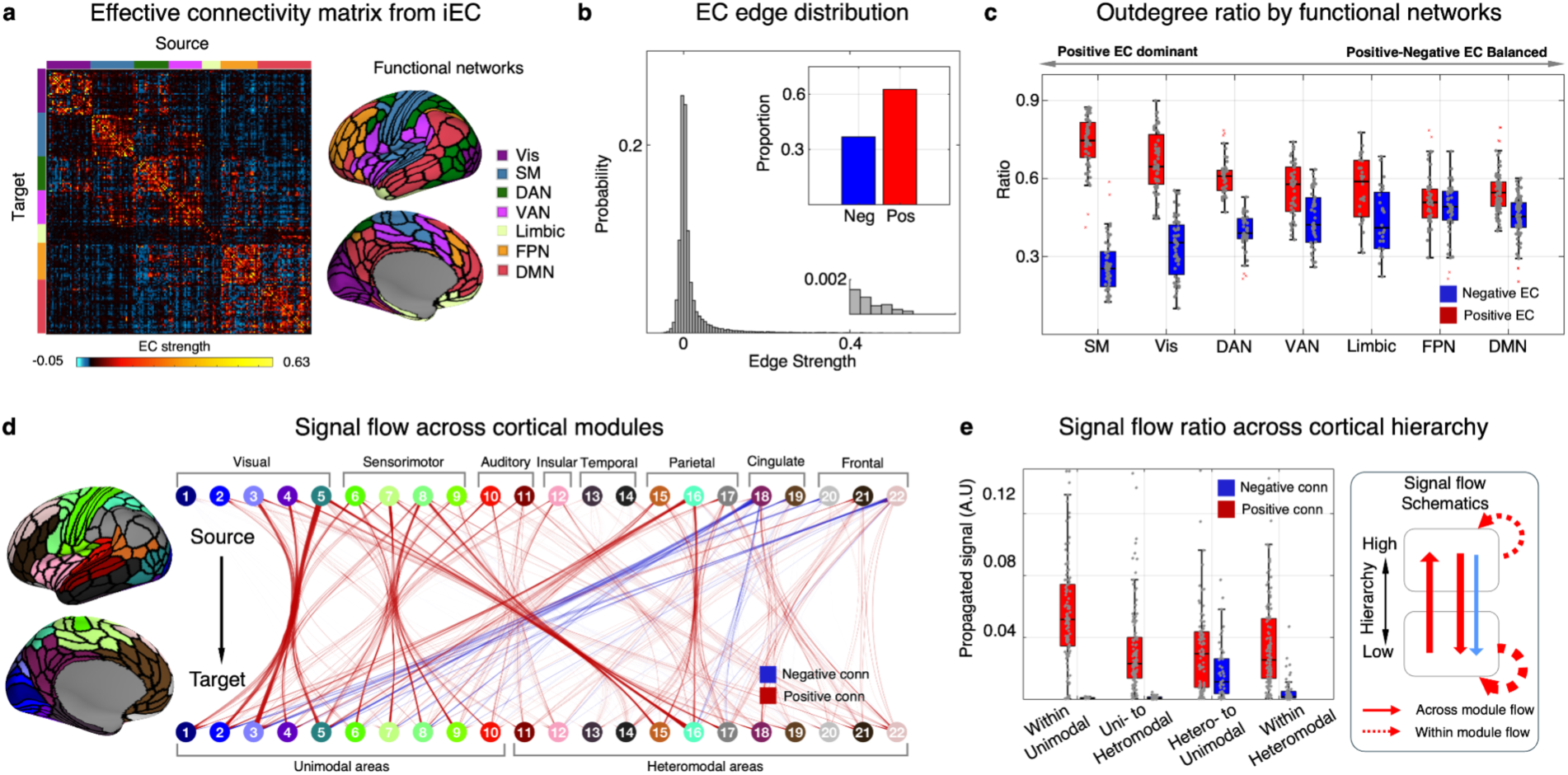
The iEC profiling of the resting-state human brain. **a**, The modular structure of iEC is unveiled by organizing it according to the 7 different functional networks of Yeo-Krienen atlas^96^ (the left cortex presented only for visualization purposes). **b**, The edge distribution of iEC shows a characteristic heavy-tail distribution. **c**, Profiling of the positive-negative ratio of outdegree connections (see **Supplementary Fig. 9c** for the indegree plot) based on the 7 functional networks. Note that the dichotomized pattern between positive and negative connections in the unimodal regions gradually converges into a form with a more balanced ratio in higher-order heteromodal areas. **d**, Signal propagation based on a linear dynamical system^18^ across 22 modules of the MMP-360 atlas. Modules are categorized into functionally specialized subnetworks for easier interpretation of the signal flow patterns. Edge thickness indicates signal strength, and color represents positive (red) and negative (blue) signal flows, respectively. **e**, Stratification of signal propagation based on the proportion of positive and negative signal flows within and between unimodal and heteromodal systems. Positive signal flow prevails within and between the systems, whereas negative signal flow is primarily observed in hetero-to-unimodal pathways during resting-state fMRI.

Next, we examined the directionality of iEC with respect to two network indices: weighted degree distribution and intrinsic signal flow. For the degree distribution, we separately estimated positive and negative iECs to check their ‘connection polarity’ (see **Fig. 4c** for outdegree and **Supplementary Fig. 9** for in-degree ECs). This revealed a discernible dominance of positive outdegree ratios within the sensory cortices, contrasted by a prominent presence of negative outdegree ratios in the heteromodal areas. Specifically, when we categorized them based on a previously established network-level hierarchy^96^, the proportion of signed connections showed a similar trend with the macaque tract-tracing result: a disproportionately large amount of positive ECs from early sensory areas and a balanced presence of positive and negative ECs from heteromodal areas, which may capture the nature of FF and FB pathways, respectively^98^.

To investigate how the estimated connectivity structure shapes the directional spread of the signals across the cortex, we have used a recently developed linear dynamical system approach^18^ to model unconstrained signal propagation from predefined seed areas (**Fig. 4d**; see **Methods** ‘*5.2 Signal flow analysis*’). This analysis revealed that positive signals predominantly originate from hierarchically lower cortical areas, whereas negative signals are almost exclusively sent from the high-order to low-level regions. Moreover, each of unimodal and heteromodal areas showed strong positive signals within their module, suggesting highly specialized functional subsystems (**Fig. 4e**). In sum, our findings show that the iEC has high potential to recapitulate the principle of signal flows across the areas with varying hierarchical levels (see **Supplementary Fig. 10** for reproducibility in the replication dataset).

#### 2.2 Retrieving directed functional hierarchy from iEC

The iEC analyses conducted so far have provided converging evidence indicating distinct signatures of directed functional connectivity that may capture a silhouette of hierarchical brain organization, including *i)* a significant relationship between iEC patterns and FF/FB connectivity in the macaque brain (**Fig. 2e**), *ii)* a different proportion of positive/negative signaling along the sensory-association axis in the human brain (**Figs. 4c** and **4d**), and *iii)* the observation that the majority of negative signal flows occurred in the heteromodal-to-unimodal pathways (**Fig. 4e**). These findings collectively motivated a systematic evaluation for whether iEC can provide any useful information to determine hierarchical levels of brain areas.

To answer this question, we employed an established modeling framework^14,100^, originally designed to quantify a hierarchical level of cortical areas and examined their presumed top-down and bottom-up connectivity topologies. Traditionally, this framework has been reserved for macaque studies due to the necessity of histological data (*e.g.,* SLN values), through which the feedforward and feedback properties of the connections could be inferred. However, as our findings (**Fig. 5a**) also provide insights into inferring the hierarchical levels of cortical areas, we used the group-level iEC matrix as a substitute for SLN to reconstruct a signal flow hierarchy map of the human brain using this framework (see **Methods** ‘*5.3 Hierarchy estimation*’).

**Figure 5.**
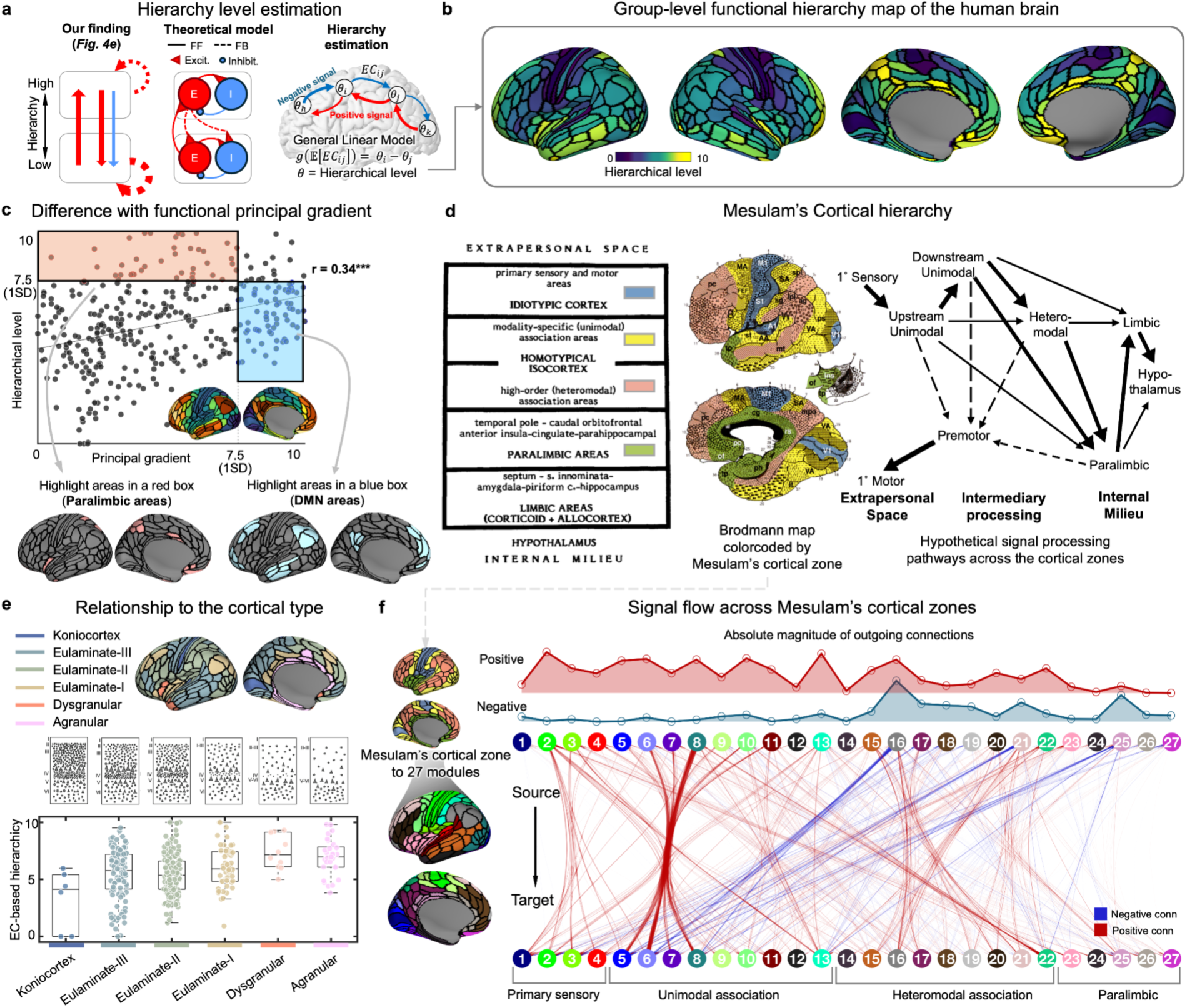
The iEC-derived functional hierarchy in the human brain. **a**, Schematic of our approach to estimate the hierarchy. To illustrate the process of hierarchy determination, we put in side by side a hierarchical signal flow from our previous findings (Fig. 4e) and its theoretical circuit model. Inspired by FLN-based hierarchy determination in the macaque brain, we used the iEC matrix to infer the hierarchy level of each cortical area across the entire human brain. **b**, The resultant functional hierarchy map is shown. The insular and paralimbic areas are positioned at the top of the hierarchy, while the primary sensorimotor areas are located at the bottom. **c**, The difference between our EC-based hierarchy and FC-based functional gradient. The brain areas with higher EC-based hierarchy (>mean+1SD) but lower functional gradient (<mean-1SD) are marked by red, while those with opposite profiles are marked by blue (*i.e.,* higher gradient but lower hierarchy) in both graphs and cortical surfaces. **d**, The cortical zones proposed by Mesulam^99^ (left), indicating different levels of cortical hierarchy. Note that within the neocortex, the paralimbic areas are at the top of the hierarchy. These four zones are superimposed on the Brodmann parcellation scheme (middle). Finally, the hypothetical signal processing pathways across corresponding hierarchical levels proposed by Mesulam are shown (right). All figures are sourced from Mesulam, 1998^99^. **e,** The relationship between the identified functional hierarchy and histological cortical types. Note that as the hierarchical value increases, the cortical type transitions from a laminar structure with clearly defined granular layers to one with less distinct or even absent granular layers. **f**, Signal propagation was analyzed across 27 modules derived from Mesulam’s cortical zones. Positive signals dominantly originated from primary and unimodal areas decreased along the hierarchy, while the negative signals progressively occupy the emission of the higher order areas.

The mapping of this signal flow hierarchy revealed a distinct and organized structure (**Fig. 5b**): primary sensory and motor areas occupy the lowest tiers, while paralimbic cortices, such as the anterior cingulate, insular, and parahippocampal regions, are positioned at the highest levels of cortical hierarchy. This pattern was partly comparable yet still distinct enough from a well-known functional gradient^16^, a dimension-reduced topographic map of undirected FC representing a sensory-association axis^29^ (**Fig. 5c**). Indeed, there was a significant spatial correlation between our EC-based hierarchy and the functional gradient map (r=0.34, p<0.001). Yet, the paralimbic regions exhibited higher values exclusively in the EC-derived hierarchy map, not int the gradient map. This emphasis on the paralimbic regions aligns with Mesulam’s initial proposition^19^ where they have been conceptualized as a ‘neural bridge’ linking the neocortex and hypothalamus (which directly monitors the internal milieu) (**Fig. 5d**).

The validity of our EC-based hierarchy map was further supported by the *post-hoc* cyto- and myelo-architecture analysis (**Supplementary Fig. 11a**). For the cytoarchitectonics, we used Campbell’s atlas^101,102^, a historical neuroanatomy resource for microscopic cellular observation. When the brain areas in this atlas were sorted based on our EC hierarchical values, the pattern closely matched a known hierarchy stream: primary sensory areas at the base, followed by unimodal and heteromodal association areas and ending with paralimbic areas at the top. A significant correlation with myeloarchitecture (by Nieuwenhuys’ atlas depicting a cortical myelin level^12^) was also found (r=0.42, p<0.001), underscoring the relationship between the signal flow hierarchy and its anatomical substrates (myelin degree), which has been previously reported to reflect the structural hierarchy of the brain.^103^.

Most importantly, our validation drew upon the seminal ‘*Structural Model*’ theory^104,105^, which posits that FF/FB connections depend on the laminar structure of the connected areas. According to this model, FB connections typically originate from areas with simpler laminar structures (‘agranular’ or ‘dysgranular’) and target areas with more elaborate laminar structures (‘eulaminate’ or ‘koniocortex’), and *vice versa* for the FF connections (*i.e.,* more elaborate to simpler laminar targeting). Inspired by this theory, we tested our hierarchy map by examining its whole-brain profile across different cortical types (**Fig. 5e**). This analysis revealed a highly significant alignment (by a general linear model; t=5.14, p<0.001), with a progressive increase in hierarchy corresponding to decrease of granular layers (see **Supplementary Fig. 11b** for the result when measured based on the functional gradient).

Finally, we again performed the unconstrained signal flow mapping on iEC, yet this time with the cortical areas sorted out by Mesulam’s cortical zones (*i.e.,* primary sensory [4 modules], unimodal association [9 modules], heteromodal association [9 modules], paralimbic [4 modules] zones). This analysis revealed that the pattern we have previously hypothesized in terms of FF (*i.e.,* largely excitatory) and FB signaling (balanced excitatory and inhibitory influences, with a latter more dominant) are well captured in our EC-based cortical hierarchy (**Fig. 5f**). Indeed, both primary sensory and unimodal association areas mostly emit positive signals, while the heteromodal regions show a balance between positive and negative signal flows, and this pattern even converts in the paralimbic region to the one with negative signals exclusively detected.

#### 2.3. State-dependent reorganization of functional hierarchy

Although so far the hierarchy derived from a resting state showed compelling biological evidence, it represents only a single brain state, one characterized by the absence of explicit environmental engagement. Therefore, whether this hierarchical structure stays in the same form no matter which brain condition is, or if it is dynamically reconfigured to adapt to contextual demands (*i.e.,* state-dependent functional reconfiguration) is still an open question.

To investigate this, we utilized two additional fMRI datasets representing major non-resting brain states. The first was acquired while participants watched a movie (’Forrest Gump’), and the second was scanned during the experience of tonic pain induced by oral capsaicin (see **Methods**, ‘*1. Data acquisition*’). The former dataset captures an externally-focused, or exteroceptive, state, characterized by attention directed toward audiovisual stimuli. In contrast, the latter represents an internally-focused, or interoceptive, state, in which awareness is concentrated on a sustained painful sensation. The functional hierarchy of each brain state was constructed using the same procedure as in our previous analyses.

When directly correlating the spatial pattern of whole-brain hierarchy, its global trend was largely preserved in both movie-watching and tonic pain states compared to the resting state (**Fig. 6a**; r=0.46/0.64, respectively; both p<0.001). Yet, scrutinzing the details of each state displayed its idiosyncratic hierarchical changes. For instance, consistent with our previous analyses (**Figs. 5e-f**), the resting state showed a monotonically increasing hierarchy across the cortical zones (**Fig. 6b**, orange). In contrast, during the movie-watching state (**Fig. 6b**, pale green), the primary sensory and unimodal association areas showed an elevated hierarchy, while the heteromodal association and paralimbic areas exhibited an opposite pattern, overall suggesting a signature of ‘*flattened cortical hierarchy*’, as also reported in a recent naturalistic fMRI study^90^. This change was explained by the subsequent signal-flow analyses (**Fig. 6c** left), in which we observed 1) an overall decrease of positive signals from the primary sensory areas, 2) reduced negative signals from paralimbic areas, and 3) an increase of negative signals in the unimodal association areas (*i.e.,* higher-order visual cortices) (all p<0.05, FDR-corrected), collectively resulting in less differentiated hierarchical levels across the whole brain. Notably, an increase of negative connections from the higher-order visual areas was found to mostly target auditory regions, potentially suggesting a competing modulation effect of shared attentional resources^106^ and/or multisensory integration^107^. This state-dependent reorganization during movie watching therefore suggests a shift in cognitive load towards externally oriented processes, as indicative of altered FF/FB signaling along sensory and attention pathways^108–110^.

**Figure 6.**
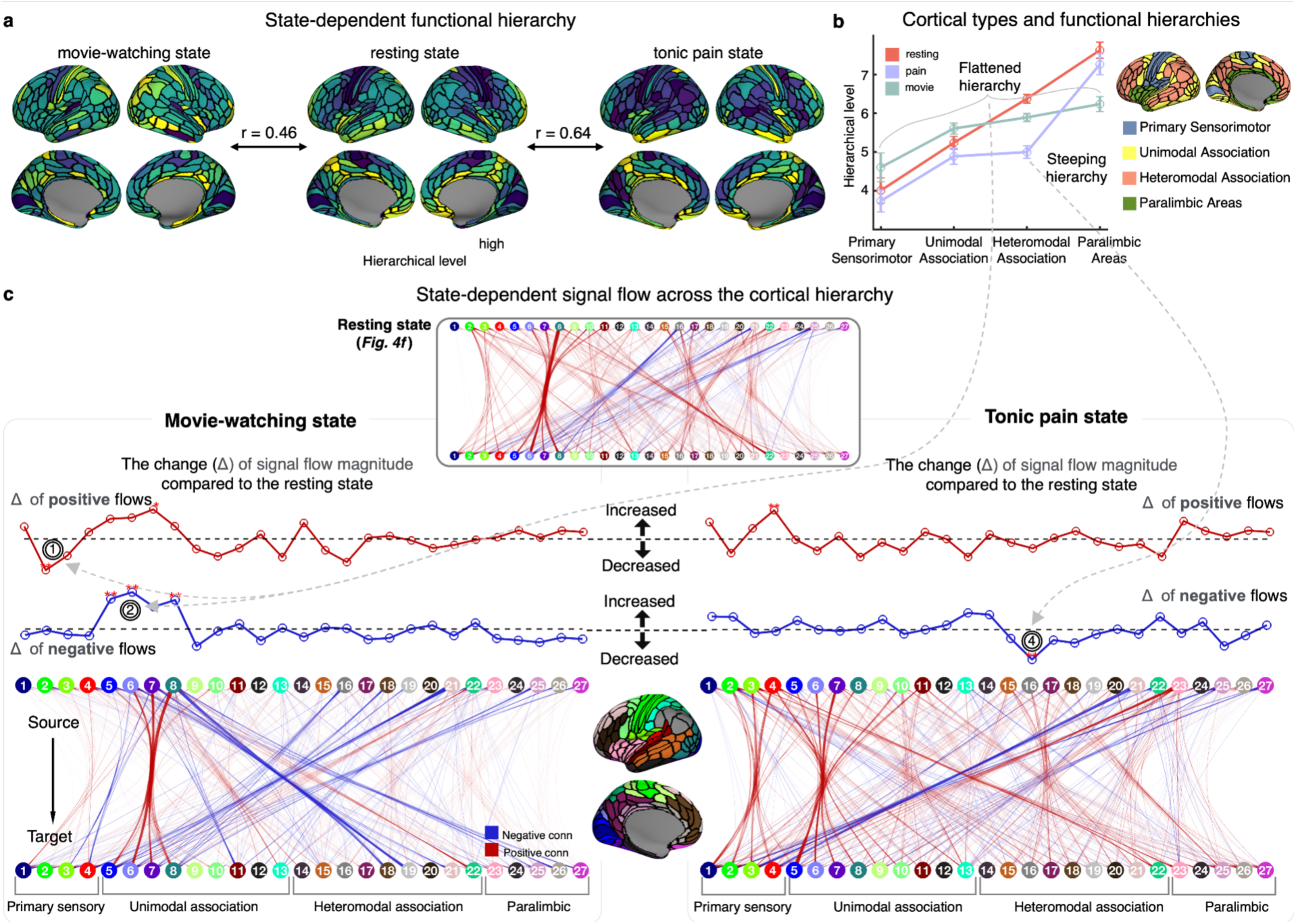
State-dependent reorganization of functional hierarchy and signal flow of the human brain. **a**, Functional hierarchy maps derived from movie-watching (left), resting (middle) and tonic pain (right) states. The correlations of their spatial patterns are shown between the states. **b**, State-dependent changes in hierarchy levels sorted out based on the Mesulam’s four cortical zones (*i.e.,* primary sensorimotor, unimodal association, heteromodal association, and paralimbic areas). Note that the movie-watching state resulted in a flattened cortical hierarchy, while the tonic pain condition accentuated the disparity between paralimbic and non-paralimbic areas (steeping hierarchy). **c**, Signal flow analysis across 27 modules of MMP (parcellated according to the cortical zones) in the movie watching and tonic pain states. (Upper) The magnitude of changes in signal flows (Δ) in these states compared to the resting state were presented as a line graph, separately for positive and negative streams and statistically tested (***: p_FDR_< 0.05, **: p<0.025, *: p< 0.05). In the graph, the findings of ⓵ ‘decreased positive signal flow’ and ⓶ ‘increased negative signal flow’ are both related to enhanced hierarchy at the primary sensory and visual association areas, while ⓷ ‘decreased negative signal flow’ implicates diminished hierarchy of the paralimbic regions, which collectively induce the whole-brain hierarchy to flatten during a movie-watching state. Similarly, ⓸ ‘decreased negative signal flow’ in the default mode area means its diminished hierarchy during a tonic-pain condition, which may cause the steeping hierarchy in this brain state. (Bottom) The edge-bundling visualization presenting overall signal flows qualitatively supports these statistical results.

In contrast, in the sustained pain condition we found an opposite pattern of hierarchical changes, that is, a ‘steeper’ hierarchy formed between the paralimbic and non-paralimbic areas (**Fig. 6b**, pale purple). Again, the signal flow mapping provided a parsimonious account for this change, demonstrating a decreased negative signal emission from the heteromodal association areas (particularly from the angular gyrus; **Fig. 6c** right), which may cause their hierarchy levels to diminish (thus resulting in a larger disparity between the heteromodal and paralimbic areas). This also indicates a state-dependent hierarchy shift, but unlike a movie-watching condition, the brain appears to focus on top-down effects of interoceptive areas by rendering the negative signals to be emitted primarily from the paralimbic system. This may suggest the prioritization of an allostatic control to cope with a potential danger from sustained pain^111,112^.

We finally note that there is also a state-general component across all three brain conditions: strong negative signal flows from ‘paralimbic/heteromodal to unimodal’ regions (especially from the cingulate and orbitofrontal areas; **Fig. 6a**), highlighting the importance of their state-invariant modulation/suppression effect. In sum, these results suggest that while the global pattern of functional hierarchy may be preserved to some extent across different states, the degree of hierarchy and its detailed functional dynamics can be significantly adjusted to more efficiently respond to the given (either external or internal) environmental changes.

## Discussion

Ever since Hubel and Wiesel’s initial discovery in the visual cortex^113^, functional hierarchy has been a central topic in systems neuroscience, serving as a primary basis to study the principle of neural information flow. While early work tended to focus on the sensory hierarchy^114,115^, the notion of this hierarchical architecture has been more broadly expanded over the last two decades, recently emphasizing the significance of its whole-brain representation spanning from the sensorimotor to association areas^9,10,18,84,103,116,117^. In this study, we reconstructed a functional processing hierarchy of the human cerebral cortex in a fully data-driven manner by analyzing different brain states across multiple fMRI datasets. Given that the hierarchy is inherently organized by ‘*directed*’ feedforward (FF) and feedback (FB) pathways^14,33^, we employed macroscale effective connectivity, linearly combining the results of existing individual algorithms (iEC). This collective inference was demonstrated not only to boost the accuracy of directionality estimation but also to allow for discovery of interesting relationships between signed (positive and negative) connections and histologically derived FF/FB pathways, providing a crucial hint to infer a hierarchical structure from in-vivo fMRI. Indeed, our iEC-based hierarchy revealed a strong alignment with a spatial distribution of hierarchy-indexed cyto- and myelo-architectonic features as well as the laminar proportion of granular cells across the brain, effectively recovering the ‘sensorimotor-association-paralimbic’ axis, as originally proposed in Mesulam’s cortical zone. Moreover, we found that this functional axis does not always adhere to the same form of hierarchy as observed in the resting state, but instead reorganizes dynamically according to brain conditions, potentially enhancing behavioral adaptability in response to rapidly changing environments. Our findings offer a novel avenue to quantitatively probe state-dependent neural information flows along the biologically validated functional hierarchy in both typical and atypical human brain conditions.

The idea behind how fMRI-based iEC enables the estimation of functional hierarchy was built upon multiple observations on our iEC profiles and their interpretation in the context of previous literature. Traditionally, the FB connections, for instance those observed between the cortex and thalamus, were primarily considered to exert a modulatory (often inhibitory) effect, refining bottom-up sensory information^118–123^. However, emerging evidence indicates a more active role of FB connections, not only as a modulator but also as a driver to increase the neural activity (excitatory)^124,125^, especially for hierarchically proximate cortical areas^85–87^. In contrast to such mixed neurophysiological effects of FB, the characteristic of FF connections has been almost exclusively associated with excitatory effects on the targeted area (but see the recent perspectives^126,127^). For example, in the macaque brain, experimentally silencing V1 through cooling strongly suppresses neural activities across the series of hierarchically connected areas from V2 to V5/MT, highlighting a significant driving role of FF pathways^128–133^.

Importantly, these distinct excitatory-inhibitory patterns between FF and FB connections also seem observable in our macroscale iEC profiles, assuming that positive iEC generally represents an excitatory effect whereas negative iEC relates more to an inhibitory effect. Parallel to this notion, when associating the sign of iEC and the patterns of supragranular labeled neurons (SLN), a histological metric to quantitatively determine the degree of FF/FB pathways in the macaque brain, we found a unique pattern of ‘*signed iEC asymmetry*’ gradient: FF connections, as determined by SLN, were characterized by a disproportionate ratio of positive iECs, whereas FB connections additionally exhibited a significant increase of negative iECs, resulting in a more balanced distribution of signed iECs. A similar asymmetry gradient was also observed in the human brain, where low-level sensory areas, known as key sources of FF signaling^1,100,129,134^, exhibited a dominant proportion of positive iECs, whereas the higher-order association areas (major sources of FB signaling^135–140)^ displayed progressively enhanced negative iECs in addition. This cross-species convergence, together with the previously hypothesized role of FF/FB connections, collectively suggest that despite its macroscale nature derived from fMRI, iEC may contain biologically meaningful signals to infer the trace of FF/FB connections, which provides a testable ground to reconstruct the functional hierarchy *in vivo*.

Given this motivation, we indeed analyzed the iEC matrix in more depth to quantitatively reconstruct the functional hierarchy, inspired by the seminal macaque studies^14,100^. Specifically, we used a general linear model (GLM) to estimate a hierarchy difference between the two brain areas (*i.e.,* source and target) according to the following, very simple rule: if the source exerts dominantly positive (or negative) influences on the target, the former cortex ranks lower (higher) than the latter one in the hierarchical axis. However, when it comes to more complex networks such as the one analyzed in our study (*i.e.,* 360 nodes that generates >10^5^ connections), calculating the analytical solution of GLM such that it 100% satisfies (without any violation) in terms of correctly ranking all brain areas in terms of their hierarchy is not possible, because of their complicatedly intertwined EC relationships. Instead, what the GLM solution provides is the hierarchy setting with ‘the least tension’ across brain regions. That is, although the hierarchy level is overall properly assigned across the brain, there could still be some areas positioned at a relatively higher hierarchical level, yet they may emit (not only negative iECs but also) a large amount of positive ECs to other brain regions (*e.g.,* heteromodal areas in our map). It should be noted, therefore, that this hierarchy assignment is not simply a problem between the two brain nodes but is a collective result, for which one has to take into account all other relationships with the rest of the brain.

Despite such computational complexity, our iEC-based method revealed several biologically noteworthy findings in hierarchy mapping. Firstly, compared to the widely used connectome compression approach called ‘functional gradient’^16^, which places the default mode network in the apex of the hierarchy, our iEC-based map puts more emphasis on the paralimbic regions to characterize a hierarchical stream of the human brain, which is in parallel to the original proposition of Mesulam^19,141^. This suggests that although many studies currently focus on the sensory-to-association axis, framing it as a major functional organization principle, the axis should also take the interoception-related cortical areas into consideration, treating them as yet another pivotal anchor. This finding highlights the importance of homeostatic and allostatic body regulations of the internal environment in goal-driven interactions with the external world. This is supported by our observation on the elevated paralimbic hierarchy (with flattened other areas’ hierarchy) during the tonic-pain condition (**Fig 6b**), suggesting that the brain may inherently prioritize the processing of interoceptive inputs over exteroceptive counterparts in certain contexts. Secondly, the validity of our hierarchy was also corroborated by histological architectural maps^12,101,102^, together with the cortical type analysis based on the ‘Structural Model of laminar connectivity’^11,105^. Notably, when we compared this hierarchy pattern to the one reconstructed based on only the positive iEC (thus assuming that their difference stems from the negative iECs), the paralimbic areas with a diminished laminar organization (*i.e.,* dysgranular and agranular cortices) exhibited the largest hierarchy changes (**Supplementary Fig. 12**) across the entire cortical zones, suggesting a significant contribution of negative iECs in precisely capturing the properties of hierarchically higher-order cortical regions, particularly for their potential FB modulation effects.

Most importantly, our findings underscore the significance of dynamic and flexible reconfiguration of the functional hierarchy based on given brain states. This phenomenon, termed ‘*state-dependent hierarchical reorganization*’, is critical for the adaptive functioning of the brain. Indeed, despite the largely static nature of its ‘anatomical’ hierarchy, the state-dependent shifting mechanism enables transiently reconfiguring a large-scale functional architecture of the brain, allowing it to effectively respond to moment-by-moment environmental and task-related changes, thereby enhancing its overall viability. Indeed, the seminal paper by Felleman and Van Essen 1991^1^ had already conjectured the possibility of this mechanism in the following sentences: *“Another possibility is that a hierarchy exists only in a loose sense, for instance, at the level of the different cerebral lobes, but not in any precisely definable manner for individual cortical areas.”,* (Providing an analogy with human societies) *“… Some organizations have an utterly rigid hierarchy, in which every individual knows precisely his or her place within a pecking order. Others are less well defined … may be inherently fluid and context dependent, in that one person ranks above another in one particular circumstance but below the other in another circumstance.”.* We believe that our discovery of the state-dependent reorganization offers novel insights into how the brain forms such a context-dependent functional hierarchy. The finding of flexibly elevated or flattened hierarchy in specific cortical regions in response to different environmental contexts exemplifies such a phenomenon. It will offer a critical hint to understand the principles of functional brain adaptability, especially when one tries to implement biologically more detailed circuit mechanisms, such as neuromodulatory controls via cortico-subcortical loops^142,143^.

Beyond the biological implications, our findings also offer methodological insights, especially for modeling studies in a large-scale network simulation of the human brain, a research field recently garnering great attention in computational and clinical neuroscience^144–148^. While previous work targeted to study effects of biophysical parameters to simulate ‘local’ dynamics (*e.g.,* recurrent excitatory and inhibitory-to-excitatory connections)^147,149^, recent evidence increasingly indicates that the area-specific properties of long-range connections could also be critical in shaping macroscale dynamics^150–152^. In parallel to this trend, our findings also highlight the role of heterogeneous connectional topology and properties, such as the embedding of cortical hierarchy, directed signal flows, and their weighted signs, in simulating more brain-like functional dynamics. Indeed, our *post-hoc* analysis suggested that the ‘negative’ iECs is, despite their substantially weak strength, essential to capture the cortical hierarchy in paralimbic regions (**Supplementary Fig 12**). Amid ongoing debates regarding the role of macroscale inter-areal connections in shaping spatiotemporal brain activity patterns^91,153,154^, our study highlights the necessity of accounting for heterogeneity in global connectivity structures, an aspect that has been largely overlooked in prior modeling research.

Effective connectivity, pioneered by Friston in 2003^155^, has evolved to encompass concepts such as directed functional connectivity and information flow^49,90^. Despite its importance, the perspectives on EC methodologies within the field have been rather divided. On the one hand, there has been genuine appreciation for the technical advancements that have led to a proliferation of algorithms aimed at estimating directionality^34,43,45–47,58^. On the other hand, this overabundance of specialized methods—each tailored to resolve specific modeling assumptions or limitations— has made it difficult for researchers to navigate the landscape and select the most appropriate approach for their scientific questions. In this context, the development of a unified framework, rather than yet another isolated algorithm, is a critical step forward. Through the integration process, we identified two key properties that help address these challenges: (1) EC algorithms can be empirically and theoretically clustered into distinct methodological categories, and (2) algorithms within the same category tend to exhibit high redundancy, while those across categories offer complementary information. For example, algorithms grounded in dynamical systems produce dense networks with Gaussian-like edge distributions, whereas graphical model-based algorithms yield sparser networks with heavy-tailed distributions. Even within the graphical models, depending on whether the method is multivariate (*e.g.,* FASK) or pairwise (*e.g.,* LiNGAM)—an algorithmic property which considers an influence of indirect, multi-hop pathways or not, the sparsity of resultant EC matrices is differently determined. Our data-driven integration consistently distinguishes these subtle differences and selects only a few non-redundant algorithms from each category, thereby leveraging their complementary strengths. The effectiveness of this integrated approach is demonstrated by its superior accuracy across different parcellation schemes and species (human and macaque), as well as its capacity to recover both static and dynamic features of empirical brain data through whole-brain simulations. Together, our framework offers a more streamlined and cohesive pathway for effective connectivity mapping in the human brain.

Several limitations should be acknowledged. First, the BOLD signal is well known for its region-specific variability in hemodynamic response function (HRF). Yet we did not correct for it in the development of our EC framework. This decision was informed by our post-hoc analyses demonstrating that all algorithms (except for MVGC) exhibited high consistency in their EC estimation between pre- and post-deconvolution (r>0.8), indicating robustness to HRF variability. Moreover, most EC algorithms, including iEC, performed better with non-deconvolved, original BOLD signals when we compared to the ground-truth directed connectivity in the macaque brain. These findings are consistent with recent work ^81^ suggesting that HRF deconvolution may, in some cases (particularly targeting large-scale whole brain neuroimaging), degrade recoverability of neural dynamics in the system identification task. While we do not discount the theoretical relevance of HRF variability, our results suggest that its practical impact on effective connectivity estimation at ‘macroscale’ may be limited. Second, our integration framework employed a linear weighting strategy, which may miss some nonlinear interactions among constituent algorithms. Despite this, the linear approach preserves the topological properties of each connectivity matrix, is computationally efficient, and offers interpretability that facilitates broader adoption. Lastly, we excluded subcortical structures (*e.g.,* thalamus and hippocampus) in our study. The primary reason is that Barbas’ Structural Model–the theoretical basis for hierarchy construction–is cortex-centric, relying on laminar-specific FF/FB projections; thus, subcortical structures fall outside the scope of this framework. Additionally, due to the characteristic low SNR of fMRI signals in deep brain structures, we observed that iEC profiles computed over the entire brain showed almost negligible EC magnitudes in those areas. Consequently, the resulting cortical hierarchy map was virtually identical when subcortical structures were included (correlation between with and without subcortices, *r*=0.96). This observation made the current study preclude them in the iEC calculation, but future studies using higher gradient-field MRI data may scrutinize the effects in depth.

## Methods

### 1. Data acquisition

In this study, we analyzed both human and non-human primate data for simulation and empirical effective connectivity (EC) analyses. In this section, we first provide the data description and then introduce the details of their image processing steps.

#### 1.1 Macaque tract-tracing data

As a first validation dataset, we leveraged macaque tract-tracing data from Froudist-Walsh *et al.* (2021)^10^. This dataset provides a comprehensive overview of cortical connectivity through retrograde tracer injections. Specifically, the tracers injected into specific cortical targets were transported to neuron cell bodies across the brain, except the injection site. These neurons, termed labeled neurons (LNs), were quantified in source areas projecting to the target. To standardize connection strengths, we calculated the fraction of labeled neurons (FLN) by dividing the number of LNs in a source area by the total LNs, excluding the injection site. Additionally, this data also included the information of supragranular labeled neurons (SLN), which indicates the proportion of labeled neurons in 2-3 layers with respect to a total labeled neuron number in the source area. This ratio has been used to characterize the feedforward and feedback nature of the connections, based on the established observation that feedforward pathways predominantly originate from the supragranular layers, whereas feedback pathways primarily arise from the infragranular layers^1,14,156^. The higher (or lower) SLN indicates the cortex predominantly involving feedforward (or feedback) pathways.

#### 1.2 Macaque neuroimaging data and preprocessing

We also analyzed 3T resting-state fMRI data from 19 anesthetized macaques from the PRIME-DE database (UC Davis site)^78^. The fMRI was acquired using Siemens Skyra scanner with 4-channel clamshell coil, with the following parameters: 1.4-mm isotropic voxels, repetition time (TR) = 1600 ms, echo time (TE) = 24 ms, field of view (FOV) = 140mm. We employed a HCP-style preprocessing pipeline to process fMRI data^157^, which included: anatomical structure reconstruction and segmentation, normalization to a standard brain template, correction for spatial and temporal distortions, and motion correction and resampling for surface and voxel data. Additional nuisance regression was applied to the fMRI data for denoising, which included: 12 motion-related regressors (motion parameters and derivatives thereof) and 5 principal components from white matter and ventricles and their derivatives (in total 32 regressors). The resulting signal was high pass filtered at 0.008Hz to remove noise induced by slow drift.

#### 1.3 Human neuroimaging data and preprocessing

##### 1.3.1 Neuroimaging datasets

Our study included three human fMRI datasets, each scanned under different brain states: a resting state, a movie-watching state, and finally a tonic-pain state. For the resting state, we have utilized the S1200 release from Human Connectome Project (HCP)^158^. After excluding participants with excessive head motion, we analyzed the data of 440 participants (mean age 28.8 years, 245 females). For a movie-watching state, we used the resource of ‘StudyForrest’ from Sengupta, et al.^159^, excluding two participants due to excessive head motion, which resulted in a final sample of 13 participants (mean age 29.4 years, 6 females). Lastly, for the tonic-pain state the data was reused from Lee, et al.^160,161^, which included 48 participants after removing those cases with an excessive head motion (mean age 22.8 years, 21 females). The details of imaging sequence are:

1. *HCP S1200 dataset*: MRI scans were conducted using a customized 3-T Siemens Connectom Skyra scanner equipped with a standard Siemens 32-channel RF receive head coil. T1-weighted structural images were acquired with 0.7-mm isotropic voxels, a TR of 2400 ms, an TE of 2.14 ms, and an FOV of 224×224 mm.
2. *StudyForrest dataset*: In this dataset, participants engaged in a naturalistic audio-visual task by watching and listening to the movie *Forrest Gump*. The MRI data were acquired using a 3-Tesla Philips Achieva dStream scanner with a 32-channel head coil. T1-weighted images were captured with 0.67-mm isotropic voxels, a TR of 2500 ms, a TE of 5.7 ms, and an FOV of 191.8×256×256 mm.
3. *Tonic-pain dataset*: To induce tonic pain, we delivered the capsaicin-rich hot sauce onto the participants’ tongues, and participants continuously rated their pain during the run. The fMRI scan duration was 20 minutes to fully cover the entire period of sustained pain from its initiation to the complete remission. The MRI were acquired using a 3-T Siemens Prisma scanner with a 64-channel head coil at Sungkyunkwan University. T1-weighted structural images were acquired with 0.7-mm isotropic voxels, a TR of 2400 ms, an TE of 2.34 ms, and an FOV of 224×224 mm.

##### 1.3.2 fMRI preprocessing

The preprocessing of these fMRI datasets was thoroughly documented in the respective original manuscript^158–161^. In brief, we aimed to adhere as closely as possible to the HCP pipeline across all datasets to maintain consistency. The pipeline included several critical steps: correcting for spatial and gradient distortions, compensating for head motion, normalizing signal intensity, removing bias fields, aligning data with T1-weighted structural images, and conforming to the 2-mm standard Montreal Neurological Institute space. Furthermore, head motion artifacts and structured noise were mitigated using a combination of independent component analysis and FMRIB’s ICA-based X-noiseifier (ICA+FIX)^162^. However, it is important to note that this specific classifier is not universally applicable to other neuroimaging data obtained with varying MRI specifications and acquisition protocols. Therefore, for the movie-watching and tonic pain datasets, we opted to employ ICA-AROMA^163^, which operates on a conceptually similar basis to ICA+FIX but is designed for more general datasets. Indeed, the two denoising approaches are only different in the way the noise components are classified, allowing us a close comparison of fMRI data across different states. Additionally, we regressed out mean CSF and WM signals from the denoised data to further exclude non-neuronal signals from our final analysis. The resulting signal was high-pass filtered at 0.008Hz to remove noise induced by slow drift. In the parcel-level analysis, we obtained the average time series for every vertex within each region defined by the Schaefer-100 (bilaterally 100 parcels) or MMP-360 (bilaterally 360 parcels) atlas.

##### 1.3.3 Diffusion MRI and structural connectivity

To provide a high-quality structural connectivity (SC) matrix for the use of a simulation-based validation, we employed independent diffusion-weighted imaging (DWI) data of 86 young-adult subjects from the Q3 release of the HCP database^164^. The HCP DWI scans were acquired using a customized 3-T Siemens Connectom Skyra scanner with a standard Siemens 32-channel RF receive head coil, with the following parameters: 2.0-mm isotropic voxels, TR = 5520 ms, TE = 89.5 ms, FOV = 208×180 mm (The detailed preprocessing steps are thoroughly documented in the original manuscript^165^).

To estimate the fiber orientation distributions^166–168^ from the preprocessed DWI data, we employed a multi-shell, multi-tissue constrained spherical deconvolution process using MRtrix3^166^. We then used a deterministic tractography algorithm to construct the tractogram from the estimated fibers^169^, employing the algorithm’s default settings with modifications only to the maximum length (250 mm) and the number of streamlines (5 million). To enhance accuracy in the tractogram, we applied a filtering method^170^, which minimizes spurious fiber tracking. SC matrices were then generated by counting the stream lines between brain regions based on two cortical parcellations: Schaefer-100^59^ and MMP-360^60^. As a final step in constructing a group-level SC matrix, we averaged the individual SC matrices.

### 2. Effective connectivity algorithms

In this study we included a diverse set of 9 EC algorithms (**Table 1**), each previously benchmarked for their ability to estimate directionality and representing the field’s major methodological paradigms.

Rather than developing a new algorithm, our objective was to unify existing approaches into a cohesive, generalizable framework that leverages their complementary strengths (*i.e.,* integrated EC, in short iEC). A cornerstone of this framework is a novel taxonomy that classifies these algorithms into two principal families: (1) *dynamical systems* and (2) *graphical models*. This classification is substantiated not only by their distinct theoretical assumptions but also by empirical, data-driven clustering of the EC results from each algorithm (**Supplementary Fig. 1**).

#### 2.1 Dynamical systems category

Dynamical systems approaches assume that observed brain signals are generated by an underlying system of interacting components evolving over time. These models formalize interactions through differential equations and often represent system dynamics in continuous time. The underlying assumption is that directional influences are embedded in the temporal evolution of the system, allowing directionality to be inferred from how past activity shapes future dynamics. Because these models explicitly track how activity in one region evolves over time in response to others, they inherently accommodate recurrent interactions such as feedback loops and cycles. Moreover, by modeling the magnitude and sign of influence, they can represent both excitatory (positive) and inhibitory (negative) effects between regions.

##### 2.1.1 rDCM (regression Dynamic Causal Modeling)^35^

A computationally efficient implementation of DCM that infers directed interactions by fitting a Bayesian linear regression model in the frequency domain. Unlike more computationally intensive DCM variants, rDCM achieves whole-brain scalability by assuming a fixed, canonical HRF and simplifying the underlying generative model. Despite these simplifications, prior benchmarking has shown that rDCM yields results comparable to those of spectral DCM, particularly for resting-state fMRI data^171^.

##### 2.1.2 VAR (Vector Autoregression)^73^

A linear autoregressive model that estimates directional influence by quantifying how past activity in one brain region predicts future activity in another. We applied a first-order VAR model (lag=1), a choice supported by previous studies^172,173^ showing that higher-order models often yield diminishing returns in the context of fMRI’s low temporal resolution^174–176^. To reduce overfitting, we applied L2 regularization to the least-squares estimation of the VAR coefficient matrix. The regularization parameter λ was optimized using a training dataset to maximize predictive accuracy with respect to the ground-truth target (*e.g.,* FLN in the macaque data), as shown in **Supplementary Fig. 13**.

##### 2.1.3 MVGC (MultiVariate Granger Causality)^73^

This extends the VAR framework by statistically testing whether the past values of one brain region improve the prediction of another region’s future activity. Specifically, it compares a full model, which includes both the target region’s own past activity and that of the potential source region, with a restricted model that includes only the target region’s own history. A significant improvement in prediction by the full model indicates a directed influence from the source to the target region.

#### 2.2 Graphical models category

Graphical models estimate directional dependencies among brain regions by analyzing statistical relationships in the data, without assuming an underlying temporal order or generative mechanism. These methods operate primarily on contemporaneous statistical structure—either pairwise or multivariate—and aim to identify the minimal set of direct interactions that best explain the observed dependencies. Directionality is inferred using various statistical criteria, including conditional independence (*e.g.,* detecting the absence of a direct connection when two regions are independent given a third), non-Gaussianity or skewness in marginal distributions, and score-based (*e.g.,* bayesian information criterion; BIC score) searches over candidate graph structures that optimize model fit and parsimony. These principles enable the construction of sparse and interpretable directed networks even in the absence of temporal resolution.

##### 2.2.1 FASK (Fast Adjacency Skewness)^40^

A two-step algorithm that first identifies adjacencies using conditional independence testing, and then orients edges based on skewness in the joint distribution. It supports feedback loops and works efficiently on high-dimensional data.

##### 2.2.2 CCD (Cyclic Causal Discovery)^74^

A method that identifies directed connections by checking whether the relationship between two regions persists after accounting for the influence of other regions. Specifically, it tests whether two regions remain statistically dependent when conditioning on different combinations of the remaining brain areas. If the connection disappears after such conditioning, it suggests that the dependency is indirect and possibly mediated. Unlike many classical Bayes-net algorithms, CCD does not assume acyclicity, allowing for recovery of cyclic (recurrent) pathways.

##### 2.2.3 BOSS (Best Order Score Search)^34^

A score-based algorithm that searches over permutations (orderings) of variables to identify the most plausible causal graph. For each candidate ordering, it constructs a directed graph by regressing each node on its predecessors and evaluates the result using a penalized likelihood score (e.g., BIC). This method balances model fit and sparsity and accommodates both acyclic and cyclic structures.

##### 2.2.4 DirectLiNGAM (Linear Non-Gaussian Acyclic Model)^75^

The model assumes linear relations and non-Gaussian noise to uniquely identify a directed acyclic graph. It infers causal order by iteratively identifying exogenous variables and removing their influences from the data.

##### 2.2.5 GRaSP (Greedy Relaxation of the Sparsest Permutation)^76^

An algorithm that estimates sparse directed networks by iteratively modifying the graph structure (*i.e.,* adding or removing edges) while evaluating each step using a scoring function (*e.g.,* BIC). GRaSP allows for minimal cyclicity and is particularly effective in high-dimensional settings where sparse connectivity is expected.

##### 2.2.6 Patel’s τ^77^

A pairwise asymmetry-based measure that infers directionality from conditional activation probabilities. Unlike the multivariate models above, it does not recover global network structure and was applied without subsampling (explained below).

To transform the inherently binary outputs of these algorithms into more continuous and stable estimates, we employed a subsampling-based ensemble approach. Following Bühlmann and van de Geer (2011)^177^, we generated 100 graphs per algorithm by applying each model to randomly drawn subsamples (each of size N/2, sampled without replacement) from the full fMRI time series of length N. We then averaged the resulting binary graphs to obtain a mean graph, where each edge weight reflects its empirical frequency across subsamples. This procedure yields continuous-valued adjacency matrices that are more robust and suitable for integration with other EC models.

All graphical models (except Patel’s τ) were implemented using the *Tetrad* toolbox (https://github.com/cmu-phil/tetrad), which provides standardized implementations of graphical algorithms. For all Tetrad-based algorithms, we adopted a common set of hyperparameters to ensure comparability. Specifically, we used the BIC score with a penalty value of 2—a widely used configuration shown to yield favorable performance in prior work^34,40,76^. Algorithm-specific parameters were left at their default values as provided by the Tetrad developers, who conducted extensive internal benchmarking across candidate configurations. These defaults reflect either optimal or highly generalizable settings as determined in the original algorithm validation studies. Interested readers are referred to the original publication of each algorithm for details.

#### 2.3 Integrated EC framework

Finally, we performed an integration of the results from individual EC algorithms to construct the iEC. In this process, the weight of each algorithm (*i.e. β* values) was determined by Bayesian optimization to maximize the objective function *ρ*:

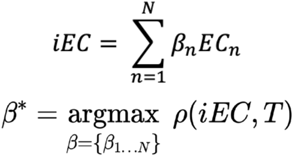

where *ρ* is the correlation between the **i*EC* and target matrix *T*. At each validation level, we employed a different target *T*: 1) At the macaque level, T was the ground-truth FLN matrix, 2) at the simulation level, it was the synthetic directed SC, and finally 3) at the human level (where ground truth is unavailable), T was defined as static and dynamical properties of empirical fMRI BOLD signals (which details are explained in the next section).

### 3. Evaluation framework and signal-level modeling

Next, to evaluate the performance of EC algorithms (*e.g.,* individual and iEC methods), we employed the following, rigorously designed, three-stage validation approach:

1. **Macaque brain validation**: ECs were estimated from resting-state fMRI of the macaque brain and compared with anatomical connectivity derived from gold-standard tract-tracing (FLN; see *1.1 Macaque tract-tracing data*). This step assessed the ability of EC methods to recover biologically validated directionality.
2. **Simulation of biologically plausible directed networks**: Second, we generated synthetic directed networks by modifying a diffusion MRI-based SC matrix. We systematically induced directionality in the SC matrix using the ‘*randmio_dir_connected.m*’ function from the Brain Connectivity Toolbox^178^. It randomly reassigns 20% of edges per realization while preserving the out-degree distribution and network connectedness. The resulting directed SC matrices retained key topological features of the empirical network, including clustering and degree distributions (pre/post rewiring correlations: r>0.9 for clustering coefficient; r>0.78 for degree distribution). These ground-truth directed networks were then used to generate oscillatory dynamics via the Hopf model. The simulated signals were in turn fed into each EC algorithm, and the resultant ECs were compared with the known ground-truth directed network.
3. **Validation using empirical fMRI signals of the human brain**: As a final validation step, ECs were inferred from resting-state fMRI of the human brain. Similarly to the previous validation, these inferred EC served as a long-range coupling matrix in the Hopf model to simulate oscillatory brain activities (see below for details). We then compared these simulated signals to empirical fMRI by computing a composite score defined as ‘Pearson correlation between empirical and simulated functional connectivity (FC)’ minus ‘Kolmogorov–Smirnov (KS) distance between their functional connectivity dynamics (FCD) distributions’ (*i.e.,* overall fit = r − KS). A higher score indicates better alignment with empirical data in terms of static and dynamical signal properties.

#### 3.1 Hopf whole-brain model

In the first and last step of our validation approach above, we simulated whole-brain signals using the Hopf model. This model combines local dynamics, described by a Landau-Stuart oscillator, with global dynamics shaped by a coupling (long-range connectivity) matrix. Despite its simplicity, the Hopf model has been widely used to simulate whole-brain dynamics and has been shown to capture numerous neurobiologically significant phenomena ^89,90,179^ (for more details, see the original paper^88^). In brief, the Hopf model is governed by the following set of coupled differential equations:

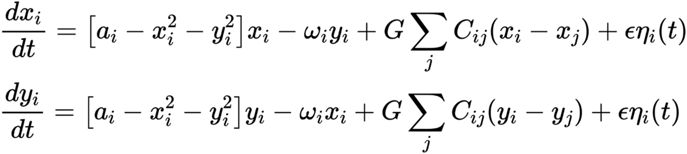

where *x_i_* and *y_i_* are state variables describing the local oscillator for region *i*, *a_i_* is a bifurcation parameter, set to −0.,01 for all regions to place them near the Hopf bifurcation point, *ω_i_* is an intrinsic frequency of each node, empirically estimated from the peak frequency of its time-series, *G* is a global scaling parameter, *C_ij_* is a connectivity matrix (either synthetic SC or EC depending on the validation stage), *η_i_*(**t**) represents additive Gaussian noise with amplitude ɛ=0.02. Simulations were performed using the Euler–Maruyama method with a time step of 0.1 to integrate the stochastic differential equations. All hyperparameter settings were adopted from prior studies^88–90^ and remained consistent with their original methodological specifications. The total simulation duration was matched to the number of time points in the empirical fMRI signal, with an initial burn-in period discarded to allow the system to reach a stable dynamical regime.

#### 3.2 Balanced Excitation-inhibition model

To evaluate the plausibility of the Hopf model as a generative framework for large-scale brain dynamics, we implemented a more biologically grounded neural mass model known as the balanced excitation-inhibition model (BEI) model^180^. This model explicitly simulates local excitatory and inhibitory populations within each brain region, governed by synaptic conductances mediated by NMDA and GABA receptors. A feedback inhibition control mechanism stabilizes local circuit dynamics by maintaining inhibitory firing rates around ∼3 Hz. Regional interactions are mediated by long-range connectivity, and simulated BOLD signals are generated using the Balloon-Windkessel hemodynamic model. We adopted all fixed parameters from the original formulation by Deco et al.^180^ and tuned free parameters to match large-scale empirical signal properties.

#### 3.3 Cross validation strategy

To ensure generalizability across our validation steps, we strictly split the dataset into a training (used for optimizing β values) and test (used for out-of-sample evaluation with the fixed β) set. Specifically, at the macaque level (N = 19), we made a half split (9 [training] vs. 10 [test]) for cross validation; at a simulation level, we repeatedly generated whole-brain signals 100 different times and split them into 20:80 training/test datasets; at the human level, we analyzed 220 HCP subjects’ data for training and another independent 220 subjects’ data for testing. Bayesian optimization was performed using MATLAB’s ‘bayesopt.m’, which constructs a probabilistic surrogate model (typically a Gaussian process) of the objective function (i.e., to maximize the accuracy of iEC) and iteratively selects coefficient values (i.e., βs of individual EC algorithms) that optimize the surrogate, balancing exploration of uncertain regions and exploitation of high-performing areas. (for more details of the algorithmic framework see Garnett 2023^181^).

#### 3.4 HRF deconvolution

Blood Oxygenation Level Dependent (BOLD) signals, the source of our EC mapping, are typically known as representing hemodynamic responses (HR) mediated by underlying neurovascular coupling. To assess the robustness of EC estimates against this HR effect and its regional variability across the whole brain, we performed a post-hoc control analysis using a data-driven HR function (HRF) deconvolution technique originally proposed by Wu et al. (2013)^182^. This method is specifically designed to remove region-specific and subject-specific HRFs from resting-state BOLD signals without explicit time information of the underlying neural events. The deconvolution approach assumes that large-amplitude spontaneous BOLD transients reflect latent neural activity and identifies these transients as pseudo-events using a point-process detection framework. For each region, an individualized HRF is then estimated by fitting a time-shifted general linear model with canonical basis functions (*e.g.,* double-gamma with derivatives), and latent neural signals are recovered via frequency-domain Wiener deconvolution. Once applying this deconvolution to the BOLD signals, we repeated the EC mapping analysis to compare the results between with and without HR deconvolution.

### 4. Neuroanatomical data

#### 4.1 Cortical types

Von Economo and Koskinas were the first to identify cortical types in the human cortex, recognizing a consistent variation of layer-specific architectonic features across different brain regions^183^. In the current study, we incorporated this cortical-type information based on the recent re-evaluation of VonEconomo’s micrographs^11^. Classification of the cortical types follows several criteria, including the development of layer IV, the prominence (marked by denser cellularity and larger neurons) of either deep (layers V–VI) or superficial (layers II–III), the definition of sublayers (such as IIIa and IIIb), the distinctness of boundaries between layers, and the presence of large pyramidal neurons in superficial layers. These cortical types reflect a spectrum of laminar complexity, from the highly elaborate *koniocortical* areas and the six distinct layers in *eulaminate III-I* to the ambiguous differentiation in *dysgranular* and the complete lack of layering in *agranular* regions. To identify the cortical types for each parcel in the MMP-360 atlas, we used the Brodmann cortical atlas along with data provided by García-Cabezas (2020^11^), which classified the cortical type of each Brodmann area. Specifically, we first assigned cortical type information to the parcels in the Brodmann atlas, and then upsampled this annotated atlas to align with the MMP-360 atlas. Additionally, we applied a general linear model to examine the linear relationship between cortical types and directed functional hierarchy levels, using ‘fitglm.m’ function in MATLAB.

#### 4.2 Cytoarchitecture and myeloarchitecture

The 1905 edition of the Campbell atlas^102^ delineates 17 distinct cortical regions, closely mirroring the cortical organization proposed by Mesulam^141^. The arealization is primarily based on histological features, such as cyto- and myelo-architectures, where regional boundaries are determined by variations in laminar arrangement and number of fibers and neurons. Each region is given a name reflective of its function and/or geographical location, examples being the visual-sensory area and the temporal area. We used data from ref^101^ for the cortical projection of the Campbell atlas in our analysis. For the myeloarchitecture data, we employed a 3D projection of the Vogt-Vogt legacy data, which is founded on a comprehensive meta-analysis and corroborated by ground-truth histological data. Detailed protocols on the myelin data projection in cortical surface can be found in the original paper^12^.

#### 4.3 Mesulam’s cortical zone

The foundational cortical zones—limbic, paralimbic, heteromodal association, unimodal association, and primary sensory-motor—were originally established in the macaque brain through extensive anatomical, physiological, and behavioral studies. Building upon this framework, Mesulam inferred the human homologues of these zones by leveraging evidence from cytoarchitectonics, electrophysiological recordings, functional imaging, and the behavioral effects of focal lesions, culminating in a complete annotation of the Brodmann areas with their corresponding cortical zones ^99^. In the present study, we translated this Brodmann-based annotation onto the MMP-360 atlas. To more effectively capture signal flow across these zones, we further refined this parcellation into 27 modules by subdividing the existing 22 modules from the MMP-360 atlas (see supplement of Glasser et al., 2016). Throughout this subdivision process, we sought to preserve the original boundaries of the MMP-22 modules wherever possible, while new modules were introduced only when a cortical zone extended beyond an existing module’s boundary, resulting in the final configuration of 27 distinct modules.

### 5. EC profiling

Below, we describe three methods that we used to profile the network characteristics of the whole-brain EC, including connectome profiles, signal flows, and the functional hierarchy estimation of the cerebral cortex.

#### 5.1 Degree and positive/negative EC ratio

For the connectome profiling, we measured both the weighted in-degree and out-degree of ECs across the whole brain’s network. This process involved calculating the number of indegree and outdegree connections for each node, providing overarching architectures across different brain regions. Given an EC matrix, with columns representing source nodes and rows representing target nodes, the weighted degree for a given node was calculated using the formula:

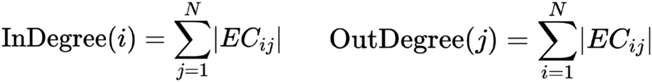

After establishing in- and out-degrees, we computed the ratio of positive to negative connections for both. This involved categorizing each connection as either positive or negative based on its sign and calculating the ratios as follows:

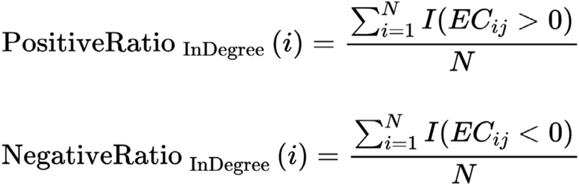

where *I* is an indicator function that yields 1 if the connection meets the given condition. The out-degree ratios were determined using a similar approach, referencing the out-degree equation mentioned above. For the statistical comparison of the degree and edge strength distributions between the target matrix (*e.g.,* synthetic directed SC in the simulation analysis) and the inferred ECs, we utilized Pearson’s correlation coefficient, implemented via the ‘corr.m’ function in MATLAB. All statistical values were FDR-corrected to account for multiple comparisons.

#### 5.2 Signal flow analysis

Beyond such graph-theoretical analyses, our EC mapping, combined with a recently proposed, linear dynamical system analysis, also provided a unique opportunity to simulate unconstrained signal propagation along the EC matrix^18^. To this end, the EC matrix was first normalized to enhance the system’s stability by adjusting its eigenvalue spectrum according to the following equation:

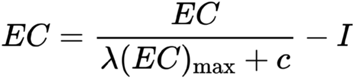

Here, *λ*is the largest eigenvalue of the EC matrix, *c* = 1 to ensure system stability, and *I* denotes the identity matrix of size N×N, which was the same as the number of parcels in the brain atlases (*e.g.,* 360 for MMP atlas). We then tracked the temporal evolution of this activity as it propagates through the network pathways, governed by the differential equation *x́* = *EC* × *x*(*t*).This equation effectively simulates the unconstrained signal flow across the network as dictated by the EC matrix. To simplify the visualization of this directed signal propagation among 360 nodes without loss of generality, we conducted this analysis within the framework of the MMP-22 modules or its finer version (MMP-27) when profiling it along the Mesulam’s hierarchy subdivision, as explained in the section *4.3*.

#### 5.3 Data-driven functional hierarchy estimation

To estimate the cortical hierarchy based on the iEC matrix, we have adopted an established model from the macaque studies^14^. This model draws upon the observation that the proportion of projections from the supragranular layers (SLN) in the source area to the target area tends to account for their hierarchical distance^1,100^. In our study, based on a series of the findings implicating the potential relationship between the signed iEC and FF/FB pathways, we substituted the SLN matrix with iEC matrix in order to calculate the hierarchy level using a following general linear model:

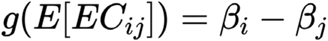

Here, *β_i_* is the hierarchical level of area *i*, and *g* is a link function that relates an expected EC_ij_ to the hierarchical distance between areas *ii* and *jj*. Extending this to the full EC matrix, we could express the model as:

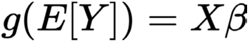

In this formulation, *Y* is 1-dimensional vectorized EC values, *β* is a vector containing the estimated hierarchical levels, and *X* is an incidence matrix (a matrix that represents which projections are connected to which brain areas) constructed from the EC, with dimensions *p* × *n*, where *p* is the number of projections and *n* is the number of areas. In our study, we followed the original approach of Barone et al. (2000)^100^ to set the link function *g* as an identity function and solved the equation using least squares. Moreover, before inputting the EC matrix, we thresholded it first by retaining the top 15% of the strongest absolute weights to obtain a balanced trade-off between sensitivity and specificity^184^. The general trend of functional hierarchy does not vary along the threshold values (see **Supplementary Fig. 14**).

#### 5.4 Assessment of state-dependent hierarchy changes

Given that so far the only target brain condition of all above-explained analyses was the resting state, we finally investigated how this hierarchy and the signal flow along its whole-brain axis could change according to different brain conditions. As specified in the previous section (*1.3.1 Neuroimaging datasets*), we analyzed the two major non-resting brain conditions, namely the externally focused movie-watching state and the pain-induced internally-oriented state. In this analysis, we first constructed the iEC-based cortical hierarchy of each state, following the exact same procedure done for the resting state. We then further quantified the change in signal flow based on the MMP-27 modules by calculating the difference between the conditions, separately for negative and positive signal flows. To evaluate statistical significance, the Δ values (*e.g.,* positive outdegree difference between movie-watching and resting states) were standardized using z-scores. Modules exhibiting statistically significant changes in signal flow were identified by applying z-score thresholds corresponding to the one-tailed (α = 0.05) and two-tailed (α = 0.025) significance levels to detect substantial deviations from the mean.

## Acknowledgements

This work was supported by the Institute for Basic Science IBS-R015-D1 (S.J.H. and C.W.W.), the National Research Foundation of Korea (NRF) grant funded by the Korea Government (MSIT) (NRF-2022R1C1C1007095, RS-2023-00217361, RS-2024-00398768 to S.J.H.), the National Science and Engineering Research Council of Canada (NSERC Discovery-1304413; B.B.), the Canadian Institutes of Health Research (FDN-154298, PJT-174995, PJT-191853; B.B.), the SickKids Foundation (NI17-039; B.B.), the Azrieli Center for Autism Research (ACAR-TACC; B.B.), BrainCanada (Future-Leaders, McConnell Brain Imaging Centre Platform; B.B.), Healthy Brains and Healthy Lives, the Tier-2 Canada Research Chairs program (B.B.), the Montreal Neurological Institute and its Centre of Excellence in Epilepsy (B.B.), Helmholtz Association’s Initiative and Networking Fund through the Helmholtz International BigBrain Analytics and Learning Laboratory (InterLabs-0015; C.P.), the Deutsche Forschungsgemeinschaft (Grant No. 491111487; C.P.), National Institutes of Health (NIH) grants RF1MH128696, R01AG047596, P50MH109429, R01MH124045, and R01MH120482 (M.M.).

