## Supplementary Materials for "*In vivo* cartography of state-dependent signal flow hierarchy in the human cerebral cortex"

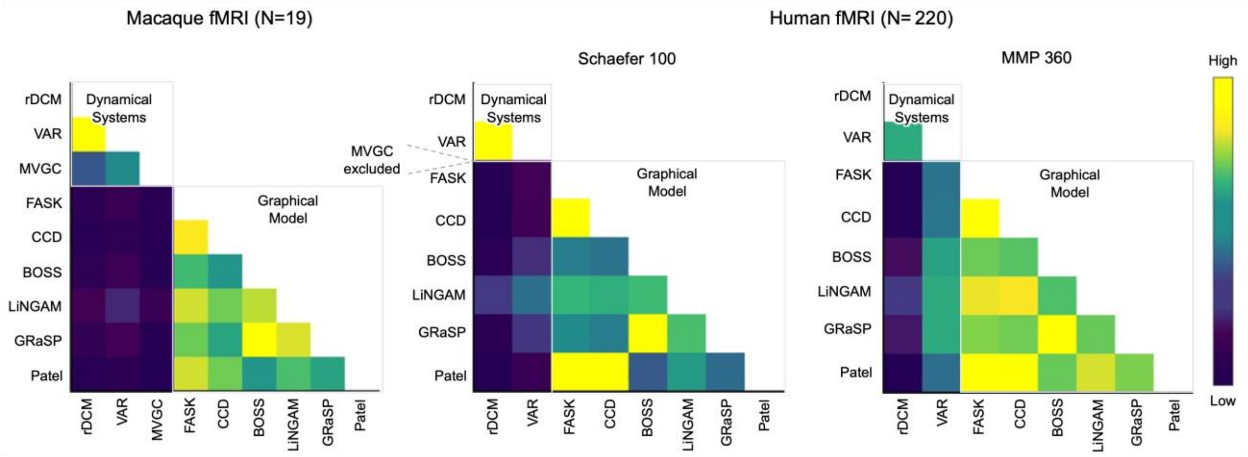


**Supplementary Figure 1. Similarity of EC patterns across algorithms in macaque and human fMRI datasets**
Pairwise Pearson correlations were computed between EC matrices derived from 9 algorithms in the macaque fMRI (N = 19) and 8 algorithms in the human fMRI (N = 220), using both Schaefer-100 and Glasser-360 parcellations. The macaque analysis included all 9 algorithms, whereas MVGC was excluded from human analyses due to its susceptibility to hemodynamic response function (HRF) variability observed in macaque data. Heatmaps indicate the degree of similarity between algorithmic outputs, with color denoting correlation strength (dart blue: low, yellow: high). Across species and parcellations, the analysis revealed consistent clustering of algorithms within theoretical families, notably among Dynamical systems–based methods (e.g., rDCM, VAR) and Graphical model–based methods (e.g., FASK, CCD, LiNGAM), providing empirical support for the proposed taxonomy of EC algorithms

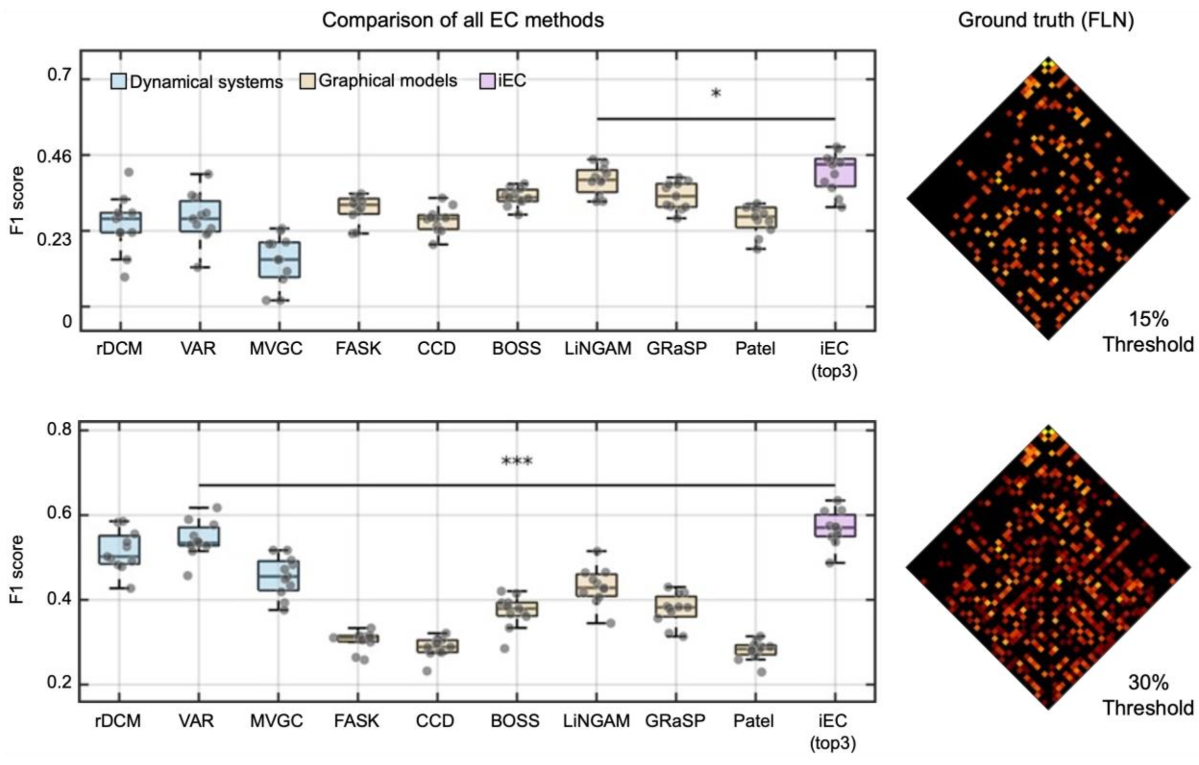


**Supplementary Figure 2. F1 scores of EC algorithms**

Both the EC and ground-truth FLN matrices were thresholded at top 15% and 30% based on the proportion of absolute connection strengths, and were fed into the computation of F1 scores. Dynamical-systems algorithms exhibited a lower F1 for a more stringent threshold (i.e., 15%), whereas Graphical-model algorithms showed an opposite pattern. At both thresholds, the iEC attained a significantly higher F1 score compared to the top individual algorithms, showing a median value of 0.56 (30% threshold, $p<0.001$) and 0.51 (15% threshold, $p=0.01$), respectively.

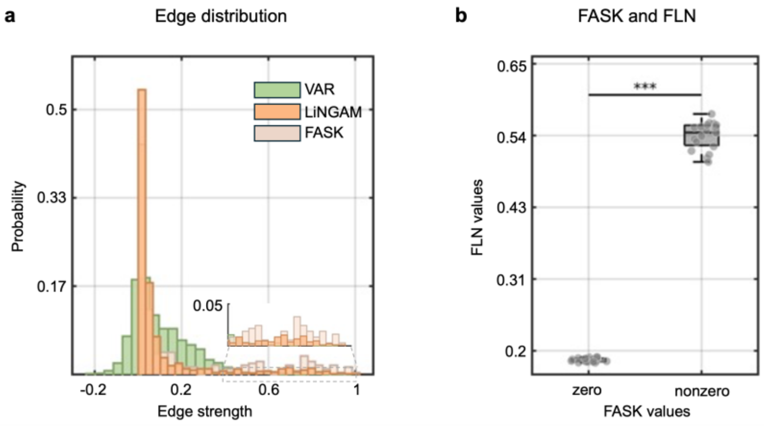


**Supplementary Figure 3. Distinct output profiles of EC algorithms**

**a**. Distribution of edge weights across three representative EC algorithms highlights qualitative differences in their output structure. The dynamical systems-based VAR model yields a dense, approximately symmetric distribution centered near zero. In contrast, graphical model-based methods such as LiNGAM and FASK produce sparse matrices with positively skewed, heavy-tailed distributions, reflecting their distinct inferential assumptions and model constraints. **b**, Comparison of macaque FLN values for edges classified as present (nonzero) versus absent (zero) by FASK. Nonzero FASK connections exhibit significantly higher FLN values (***$p < 0.001$, Wilcoxon rank-sum test), suggesting that FASK preferentially recovers sparse but biologically potent edges.

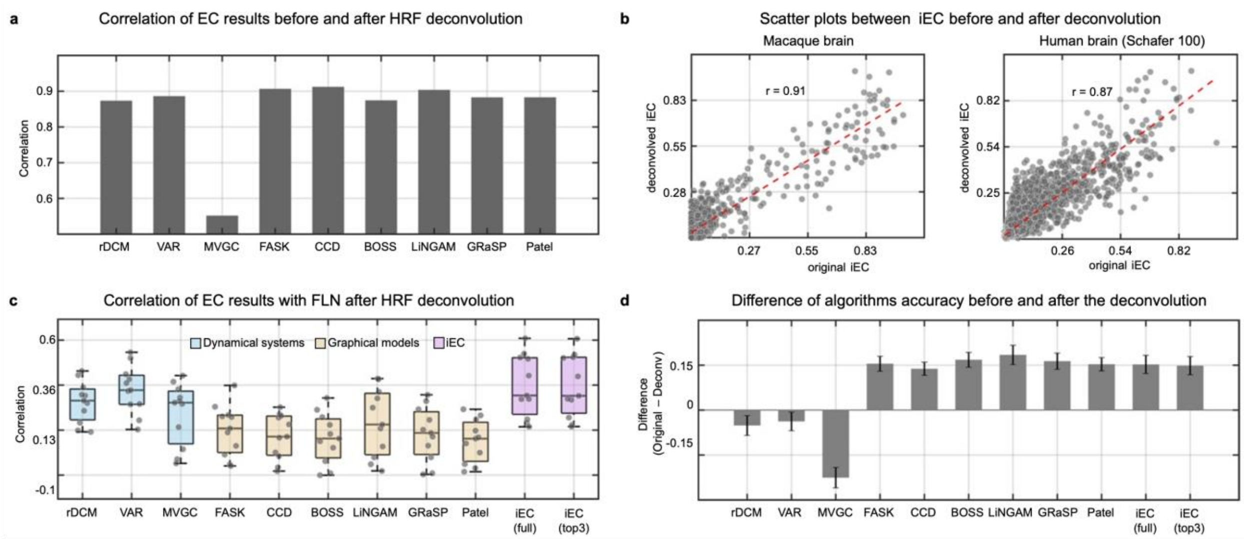


**Supplementary Figure 4. The impact of HRF deconvolution on the EC estimates**

To evaluate the influence of hemodynamic response function (HRF) variability on EC estimation, we applied a blind deconvolution approach18, designed to remove region-specific HRFs from observed BOLD signals. **a**, Group-level correlations between EC estimates obtained before and after HRF deconvolution in the macaque brain. All algorithms, with the exception of MVGC, exhibited high consistency (r > 0.8), suggesting that most methods are robust to HRF variability. **b**, Scatter plots comparing original and deconvolved iEC edge strengths for macaque (left) and human (Schaefer-100 parcellation; right) datasets. Each dot represents a single directed connection. The strong correlations (r = 0.91 for macaque; r = 0.87 for human) indicate that the iEC framework is stable and resilient to HRF-induced distortions. **c**, Accuracy of EC methods after HRF deconvolution, as measured by correlation with the ground-truth FLN. Each point reflects an individual subject. **d**, Differences in EC method performance before and after HRF deconvolution. Bar heights indicate the mean differences in accuracy (correlation with FLN) between the original and deconvolved EC estimates, with error bars representing the standard error of the mean (SEM). Overall, most of EC algorithms (including iEC) were either mildly affected by HRF deconvolution or demonstrated a superior accuracy in EC mapping when using the (non-HRF-deconvolving) original signals.

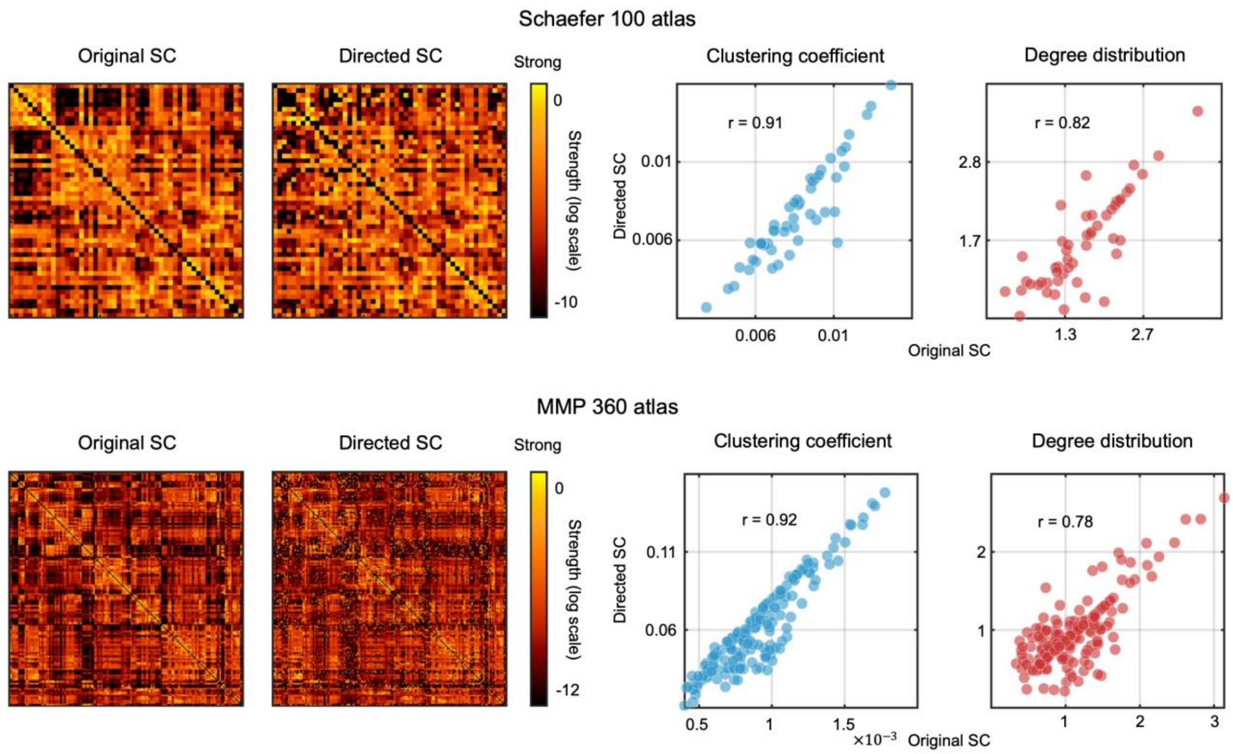


**Supplementary Figure 5. Comparison between original and synthetic (directed) SC**

Examples illustrating the construction of directed SC matrices derived from the original group-level SCs for Schaefer-100 and Glasser-360 atlases. Only the left hemisphere is displayed for simplicity, and the connection strengths were log-transformed to improve visualization: upper row (Schaefer-100), lower row (Glasser-360). Matrices depict the original SC (left) and the synthetic directed SC constructed using the 'randmio_dir_connected' algorithm (middle). The rightmost two scatter plots show the correlation of topological features (i.e., clustering coefficient and degree distribution) between the original and directed SCs. Overall, both visual inspection and correlation analysis reveal a preservation of overall network characteristics, confirming a minimal distortion of the SC network after the introduction of directional connections.

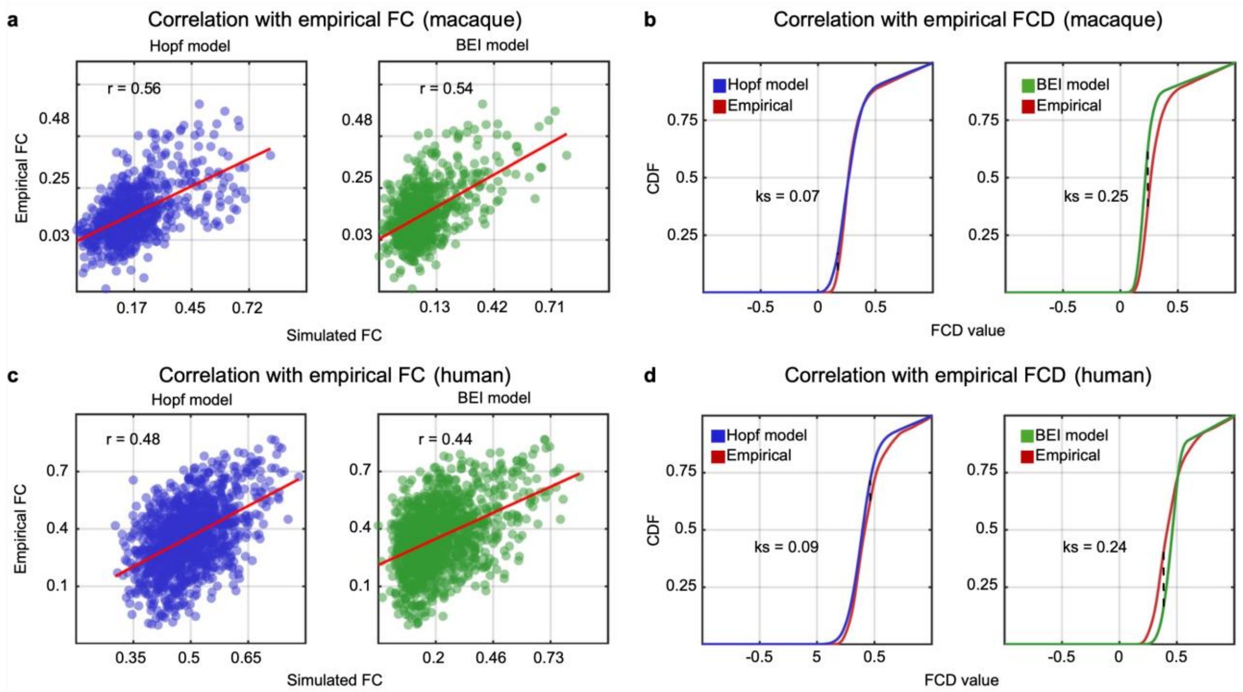


**Supplementary Figure 6. Comparison of the simulation results between the Hopf model and the biophysical model.**

The functional connectivity (FC) and functional connectivity dynamics (FCD) obtained from empirical data were compared to those generated by each simulation model. **a**, For macaque data, simulations were performed using the group-level FLN matrix as a structural backbone for an underlying directed network. Both models produced FC patterns that correlated with empirical FC, with the Hopf model showing a slightly higher correlation ($r = 0.56$) than the BEI model ($r = 0.54$), as indicated by red linear fits. **b**, FCD distributions were evaluated using cumulative distribution functions (CDFs) of the lower-triangular elements of the FCD matrices, and their similarity to empirical data was assessed using the Kolmogorov–Smirnov (KS) statistic. The Hopf model more closely matched the empirical FCD (KS = 0.07) than the BEI model (KS = 0.25), with the two model distributions differing significantly ($p = 2 \times 10^{-27}$, KS test). **c**,**d**, Same analysis as in (**a**,**b**), but applied to human fMRI (Schaefer-100). Here, the simulations were based on group-level structural connectivity restricted to the left hemisphere. The Hopf model again showed a higher correlation with empirical FC (r = 0.48 vs. 0.44) and a better match to empirical FCD distributions (KS = 0.09 vs. 0.24), with a significant difference between model FCDs ($p = 1\times10^{-24}$, KS test). These results highlight the Hopf model's sufficient validity to capture both static and dynamic features of empirical brain activities across the two primate species.

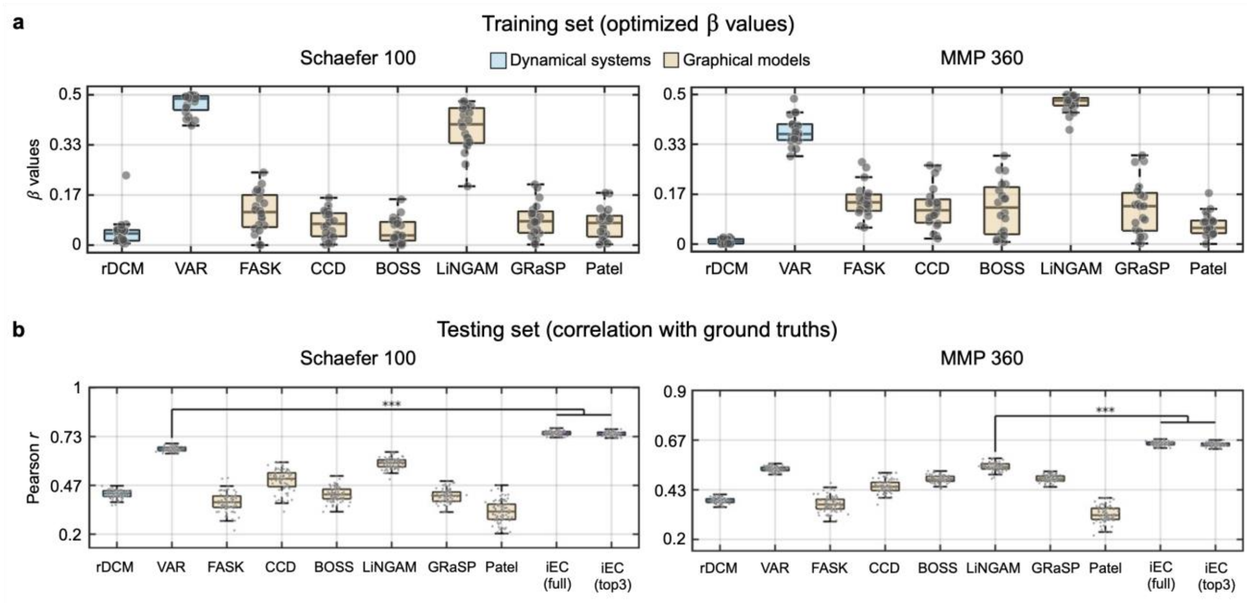


**Supplementary Figure 7. Simulation-based validation of the iEC framework**

**a**, Optimized βs derived from the training data (N=20 out of simulations using 100 synthetically generated, directed networks from human SC) across two different parcellations (Schaefer-100 and MMP-360). The distribution of βs is consistent across both network resolutions; VAR consistently emerged as the strongest algorithm within the dynamical systems category, while LiNGAM consistently demonstrated the highest β within the graphical model category. FASK ranked third in median β value for both resolutions. **b**, Correlation between the EC estimates and their underlying ground truths (directed SC matrices) obtained from the testing data (N=80). The full iEC framework (using all 8 algorithms) significantly outperformed the best-performing individual algorithm at both network resolutions (Schaefer-100: $r = 0.75$; MMP-360: $r = 0.65$; both $p < 0.001$, Wilcoxon rank-sum test). Moreover, the reduced model constructed from only the top three algorithms (VAR, FASK, LiNGAM) exhibited strikingly similar performance relative to the full iEC model, highlighting the robustness of the algorithm selection process.

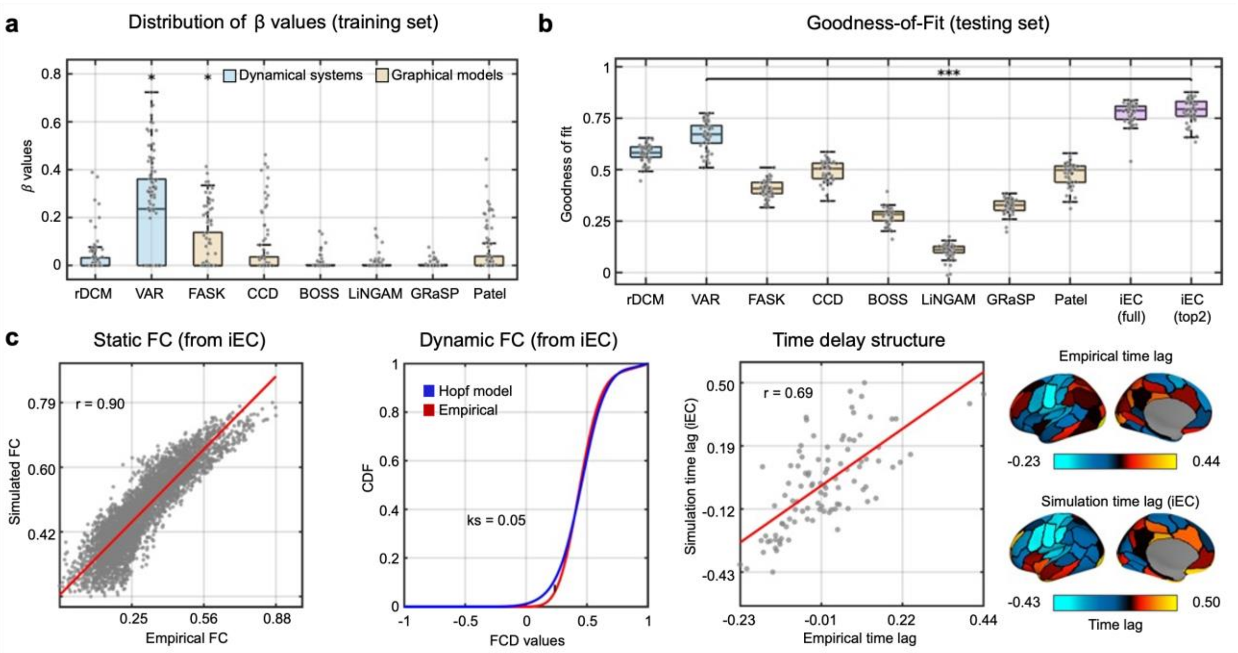


**Supplementary Figure 8. Validation of iEC framework using human fMRI data (Schaefer-100 atlas)**

**a**, Distribution of the optimized βs across 50 independent runs of Bayesian optimization on the training set. Each dot represents the optimal weight obtained from a single run, reflecting each algorithm's relative contribution to the integrated EC model. **b**, the overall fit scores of each EC algorithm and iEC evaluated in a held-out testing set (N = 220). The fit was computed as the static FC correlation minus the FCD KS distance. iEC variants (full model and top-2 contributing algorithms) exhibited superior fit relative to individual EC algorithms. Each dot represents one realization of the simulation. **c**, Comparison of empirical and iEC-simulated FC/FCD. (left) Each dot represents a FC value between a pair of brain regions. The red line shows the least-squares linear fit. (Middle) Cumulative distribution functions (CDFs) of FCD from empirical and simulated data, with the distance along the y-axis indicating a KS distance. (Right) Comparison of empirical and simulated time delay structure. Each dot in the scatter plot represents the mean time lag of a brain region; red line denotes the best linear fit. Group-averaged delay maps are shown on the cortical surface.

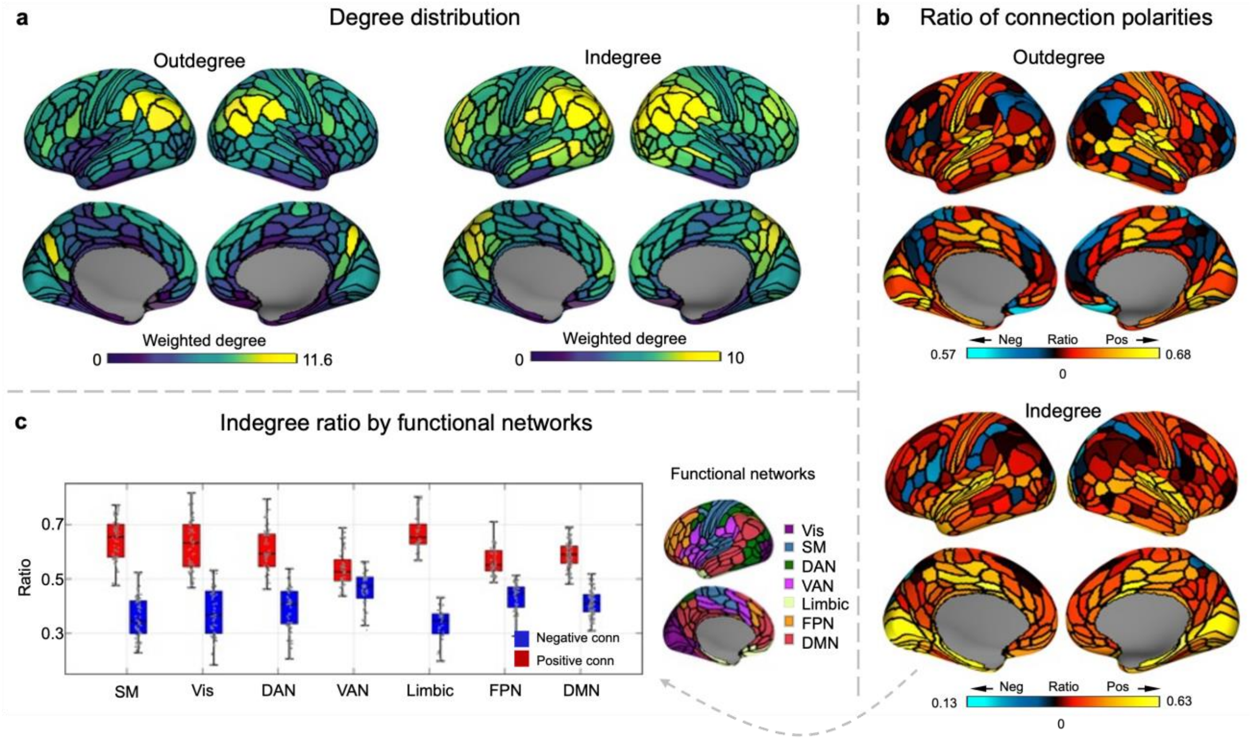


**Supplementary Figure 9. The iEC network profiling of the resting state human brain**

**a**, The degree distribution of the iEC. While the angular gyrus and precuneus show increase in both out- and in-degree, the indegree was more pronounced in a wider coverage of DMN (e.g., frontal and temporal cortex) during the resting state. **b**, The positive-negative ratio of outdegree and indegree connections is shown. (Outdegree) Heteromodal and basal forebrain regions are skewed toward negative connections, while the primary sensory areas predominantly feature positive connections. (Indegree) The dorsal and ventral attention networks receive more negative than positive connections, while the early visual area, medial temporal lobe, and insula show a higher proportion of positive connections. **c**, The distribution of positive and negative 'incoming' connections profiled according to the Yeo-Krienen atlas. The ventral attention network displays the most balanced ratio of positive to negative connections, while the limbic areas receive the fewest negative connections.

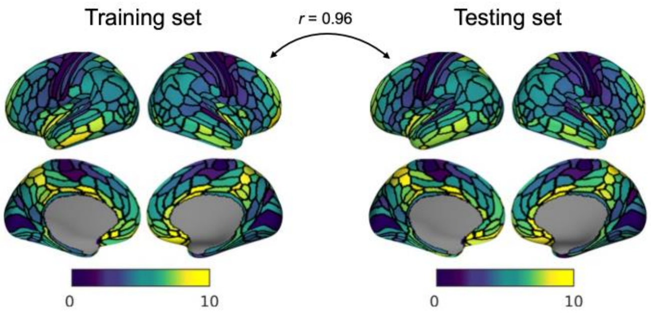


**Supplementary Figure 10. Replicability of cortical hierarchy maps across independent training and testing sets** Cortical hierarchy maps derived from iEC were independently estimated using training and testing datasets. The iEC in each map was computed based on β values obtained from the opposite set (i.e., training β values applied to testing data, and vice versa). The resulting maps showed highly similar spatial patterns, with a strong correlation between them (r = 0.96), indicating high replicability of the hierarchy structure across datasets.

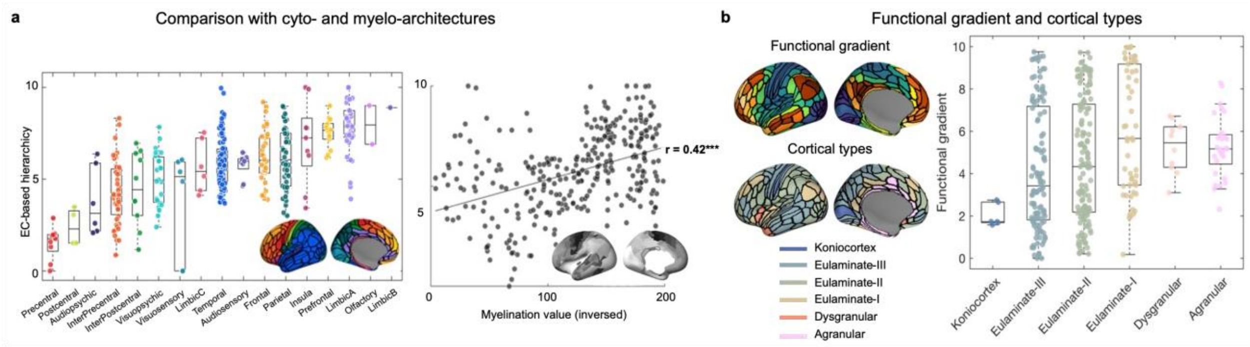


**Supplementary Figure 11. functional gradient measured using cortical type atlas**

**a**, The relationship between the identified functional hierarchy and cyto- and myelo-architectural features derived from histological atlases. **b**, Functional connectivity gradient sorted out according to hierarchically ordered cortical types. Unlike our iEC-derived functional hierarchy, which shows a gradual increase towards dysgranular and agranular areas, the gradient value showed *i)* generally larger variability across cortical types and *ii)* a peak at the eulaminate-I, which encompasses largely the default mode network regions

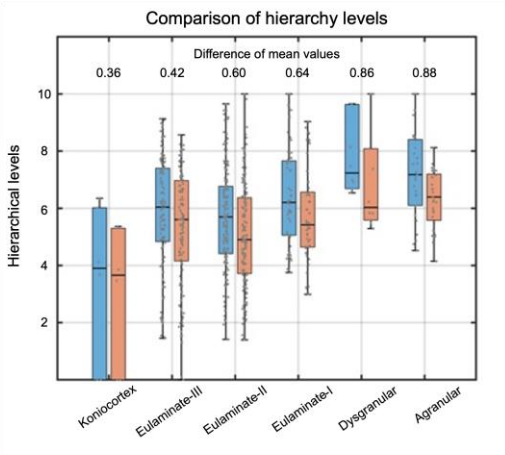


**Supplementary Figure 12. Impact of negative connections on the signal flow hierarchy**
To assess the impact of negative connections on hierarchy estimation, we compared the directed functional hierarchy derived from the original intact iEC and the one constructed by only positive iEC. Significant differences in hierarchy were observed across all cortical types except for the koniocortex. Notably, the disparity in median values increased as cortical types transitioned toward less or non-granular regions, suggesting that the inclusion of negative connections enhances hierarchy estimation in the higher-order cortical areas.

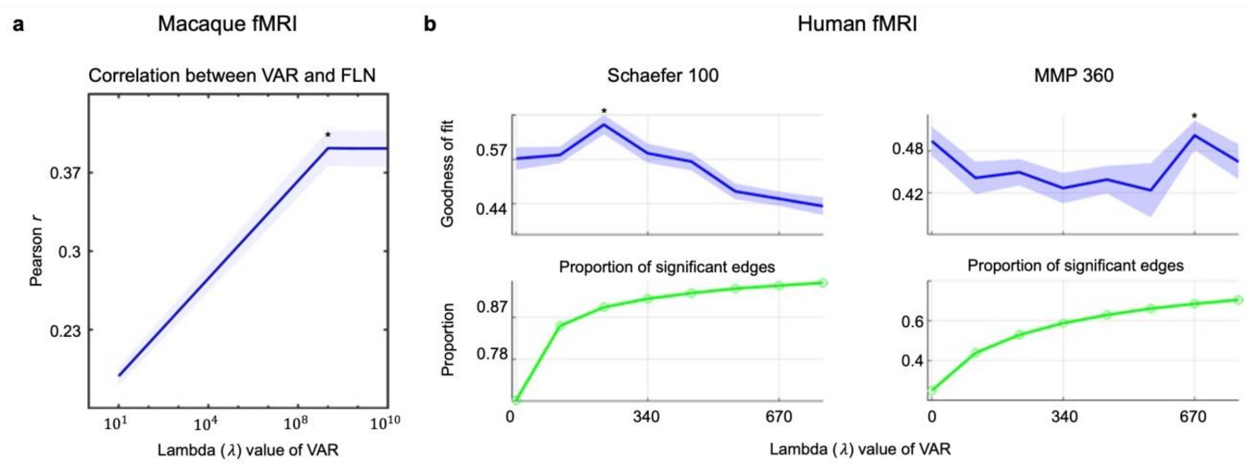


**Supplementary Figure 13. Optimization of the regularization parameter (λ) in the VAR algorithm using ground-truth and empirical data**

**a**, In the macaque fMRI dataset, the optimal λ value was selected to maximize the correlation (Pearson's r) between the estimated EC from VAR and the ground-truth anatomical connectivity (FLN). Correlation increased monotonically with λ and saturated at high values ($\lambda \geq 10^8$). **b**, In human fMRI datasets, where ground-truth EC is unavailable, λ was optimized by maximizing the overall fit between simulated and empirical brain activity, shown for both Schaefer-100 and MMP-360 atlases (top row). Asterisks indicate the λ value yielding peak overall fit. The corresponding proportion of significant edges (bottom row) increased with λ, reflecting a trade-off between model sparsity and fit.

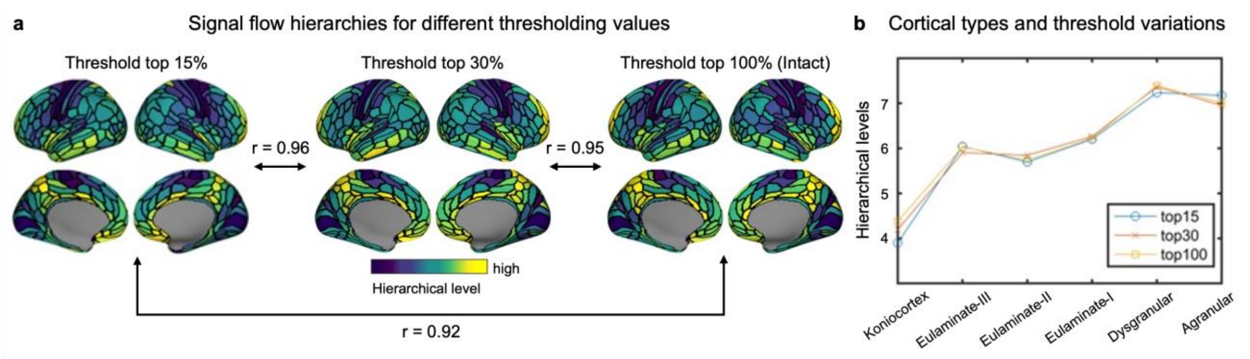


**Supplementary Figure 14. Comparison of signal flow hierarchies for different threshold values**
**a**, Signal flow hierarchy maps constructed based on the iEC thresholded by various cut-off values (15, 30, 100%). The organizational structure of the signal flow hierarchy remains robust across different thresholds, exhibiting a high correlation of >0.9 r-values between the maps. **b**, Median values for each cortical type were measured from the signal flow hierarchy maps derived from different thresholding values. The patterns show a highly significant overlap, indicating a stable and robust detection of our signal flow hierarchy against different network thresholds.
